# Navigating the amino acid sequence space between functional proteins using a deep learning framework

**DOI:** 10.1101/2020.11.09.375311

**Authors:** Tristan Bitard-Feildel

**Affiliations:** Sorbonne University

**Keywords:** Keywords, Deep learning, latent space exploration, protein sequence, protein function

## Abstract

**Motivation:** Shedding light on the relationships between protein sequences and functions is a challenging task with many implications in protein evolution, diseases understanding, and protein design. Protein sequence / function space is however hard to comprehend due to its complexity. Generative models help to decipher complex systems thanks to their abilities to learn and recreate data specificity. Applied to protein sequences, they can point out relationships between protein positions and functions capture the sequence patterns associated with functions or navigate through uncharted area of molecular evolution.

**Results:** In this study, an unsupervised generative approach based on adversarial auto-encoder (AAE) is proposed to generate and explore new sequences with respect to their functions thanks to the prior distribution allowing a continuous exploration of the latent space. AAEs are tested on three protein families known for their multiple functions. Clustering results on the encoded sequences from the latent space computed by AAEs display high level of homogeneity regarding the protein sequence functions. The study also reports and analyzes for the first time two sampling strategies based on latent space interpolation and latent space arithmetic to generate intermediate protein sequences sharing sequential and functional properties of original sequences issued from different families and functions. Generated sequences by interpolation between latent space data points demonstrate the ability of the AAE to generalize and to produce meaningful biological sequences from an evolutionary uncharted area of the biological sequence space. Finally, 3D structure models generated by comparative modelling between different combinations of structures of different sub-families and of generated sequences from latent space or sub-family sequences point out to the ability of the latent space arithmetic to successfully transfer functional properties between sub-families. All in all this study confirms the ability of deep learning frameworks to model biological complexity and bring new tools to explore amino acid sequence and functional spaces.

**Availability:** Code and data used for this study are freely available at https://github.com/T-B-F/aae4seq.

**Supplementary information:** Supplementary data are available at online.

## 1 Introduction

Protein diversity, regarding sequence, structure or function, is the result of a long evolutionary process. Protein fitness and natural selection lead to the current observation of only a fraction of all possible amino acid sequence combinations and therefore, structures and functions. These observed sequences are also referred to the amino acid sequence space. Exploring the sequence space and understanding its hidden constrains is highly challenging due to its huge size [1, 2]. The classification of amino acid sequences into protein domain families allows to organize the sequence space and reduce its complexity.

Many resources have been developed over the years to group amino acid sequences into families with members sharing sequence and structural similarities which reduce and organize the sequence space [3, 4, 5]. However, the sequence space area between these families, and even inside a family as one family may gather several groups of amino acids with different molecular functions [6], are mostly uncharted. Navigating the sequence space with respect to the functional diversity of a family is therefore a difficult task whose complexity is even increased by the low number of proteins with experimentally confirmed function. In this regard, computer models are needed to explore the relationships between sequence space and functional space of the protein families [7, 8, 9, 10, 11, 12].

Understanding the relationships between amino acid positions of a sequence groups responsible for a particular molecular function and how to cross the sequence space from one group to an other have a lot of implications in molecular engineering, functional annotation, and evolutionary biology. In this study, tools and strategies based on an unsupervised deep learning approach are proposed to model and navigate the evolutionary uncharted area of the amino acid sequence space.

Previous deep learning generative models such as variational autoencoders (VAE) have been applied on biological and chemical data to, for example, explore and classify gene expression in single-cell transcriptomics data [13] or explore the chemical space of small molecules for drug discovery and design [14, 15]. Their ability to reduce input data complexity in a latent space and to perform inference on this reduced representation make them highly suitable to model, in an unsupervised manner, complex systems. Regarding protein science, VAE have been able to accurately model amino acid sequence and functional spaces [16], predict mutational impact [17, 18], decipher protein evolution and fitness landscape [19] or design new proteins [20]. In this study, Adversarial AutoEncoder (AAE) network [21] is proposed as a new and efficient way to represent and navigate the functional space of a protein family. Unlike VAE, AAE networks constrain the latent space over a prior distribution which allows better inference to explore the whole latent distribution, which is particularly useful for travelling through uncharted area[22].

Like VAE and other autoencoder architectures, AAEs reduce high dimensional data by projection, using an encoder, to a lower dimensional space. This space is known as a latent space or embedding representation which in turn can be reconstructed by the decoder. AAE [21] architecture corresponds to a probabilistic autoencoder but with a constraint on the latent space of the encoder which follows a defined prior distribution. This constraint is applied using an generative adversarial network (GAN) [23] between the latent space and the prior distribution. It ensures that meaningful samples can be generated from anywhere in the latent space defined by the prior distribution. Applied to biological sequences of a protein domain family, it is then possible to encode the sequence diversity to any prior distribution and thus, to sample and generate new amino acid sequences of the family from any point of the prior distribution. Ideally, the learned latent space should be representative of the functions of considered the protein domain family.

The clusters induced by latent space of AAE network were analyzed to verify their ability to group sequences according to function as observed with VAE networks. Three protein families including different sub-families were used to train AAE models and to analyze the clustered sequences produced by the models using their functional annotation. The three different protein families selected were the sulfatases, the HUP (HIGH-signature proteins, UspA, and PP-ATPase) and the TPP (Thiamin diphosphate (ThDP)-binding fold, Pyr/PP domains) families. The sulfatases are a group of proteins acting on sulfated biomolecules. This family have been manually curated into sub-family with specific functions according to substrate specificity [24]. They are found in various protein family databases, such as in Pfam (PF00884). However, as mentioned previously, they can have different substrate specificity despite being in the same family. The SulfAtlas database [24] is a collection of curated sulfatases centered on the classification of their substrate specificity. The majority of Sulfatases (30,726 over 35,090 Version 1.1 September 2017) is found in family S1 and is sub-divided into 73 sub-families corresponding to different substrate specificities. Sub-families S1-0 to S1-12 possess experimentally characterized EC identifiers.

The two other protein families, HUP and TPP families are not manually curated but were selected as they are known to have multiple functions [6]. Proteins of the HUP family are a very diverge group with functions linked to particular motifs such as HIGH and KMSKS (nucleotidyl transferases and t-RNA synthetases activities), ATP PyroPhosphatase motif, or sequence motifs responsible of the hydrolysis of the alpha-beta phosphate bond of ATP [25, 26, 27]. The TPP family is made of very similar protein domains which are probably evolutionary related [28, 29]. They have pyruvate dehydrogenases, decarboxylate and binding functions [28].

VAE have often be used to cluster protein sequences and to interpret the resulting clusters regarding their function or evolutionary history but few experiments have been performed on their generative ability, particularly when it comes to navigate or to transfer features between clusters. In this study, two experiments were carried out in this direction using latent space interpolation and latent space arithmetic operations respectively. These experiments were designed as new tools and frameworks for the amino acid sequence space exploration.

The latent space coverage of the protein domain family functional space was analyzed using data point interpolations between protein sequences of different sulfatase sub-families encoded in the latent space. The interpolated data points correspond therefore to to unseen proteins, *i.e.* evolutionary uncharted area between groups of amino acid sequences. A good model should be able to produce realistic protein sequences from these data points.

To go beyond the functional clustering induced by generative models such as VAEs or AAEs, this study also explored arithmetic operations with protein sequences encoded in their latent space to generate new protein sequences. Arithmetic operations on latent space have previously been reported to transfer features between images of different classes [30] and may therefore have interesting potential for molecular design and for exploration of the amino acid sequence space. Four different strategies were explored to combine latent spaces of different sulfatase sub-families. The generated proteins from the combined latent spaces were analysed in term of sequences and structures, after being built by comparative modelling.

## 2 Methods

### 2.1 Protein Families

#### The sulfatase family

An initial seed protein multiple sequence alignment was computed from sequences of the protein structures of SulfAtlas [24] database sub-families 1 to 12. This seed was used to search for homologous sequences on the UniRef90 [31] protein sequence database using hmmsearch [32] with reporting and inclusion e-values set at 1*e*^−3^.

A label was assigned to each retrieved protein if the protein belonged to one of the 12 sub-families. The multiple sequence alignment (MSA) computed by hmmsearch was filtered to remove columns and sequences with more than 90% and 75% gap characters respectively. Proteins with multiple hits on different parts of their sequences were also merged into a single entry. From 105181 initial protein sequences retrieved by hmmsearch, the filtering steps led to a final set of 41901 proteins.

#### HUP and TPP protein families

A similar protocol was followed for the HUP and TPP protein families. Instead of using an initial seed alignment made of sequences with known 3D structures, the CATH protein domain HMM [33, 34] was used to search for homologous sequences in the UniRef90 database. CATH models 3.40.50.620 and 3.40.50.970 correspond to the HUP and TPP protein families, respectively. A sequence filtering pipeline identical to the one used for the sulfatase family was applied to each of the resulting multiple sequence alignments.

The final number of proteins in each dataset was: 25041 for the HUP family (32590 proteins before filtering) and 33693 for the TPP family (133701 before filtering).

### 2.2 Deep learning model

#### Generative Adversarial Network

A complete description of Generative Adversarial Network can be found in (author?)[23]. To summarize, the GAN framework corresponds to a min-max adversarial game between two neural networks: a generator (G) and a discriminator (D). The discriminator computes the probability that an input *x* corresponds to a real point in the data space rather than coming from a sampling of the generator. Concurrently, the generator maps samples *z* from prior *p*(*z*) to the data space with the objective to confuse the discriminator. This game between the generator and discriminator can be expressed as:

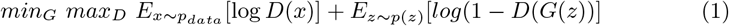

#### Adversarial auto-encoder

Adversarial autoencoders (AAEs) were introduced by (author?)[21]. The proposed model is constructed using an encoder and a decoder networks, and a GAN network to match the posterior distribution of the encoded vector with an arbitrary prior distribution. Thus, the decoder of the AAE learns from the full space of the prior distribution. The model used in this study to compute the aggregated posterior *q*(*z|x*) (the encoding distribution) uses a Gaussian prior distribution. The mean and variance of this distribution is predicted by the encoder network: *z*_*i*_ ~ *N* (*µ*_*i*_(*x*), *σ*_*i*_(*x*)). The re-parameterization trick introduced by (*author?*)[35] is used for back-propagation through the encoder network.

Three different architectures were evaluated. The general architecture is as follows (see Supplementary Table 1 and Supplementary Figure 1 for a representation of architecture number 3). The encoder comprises one or two 1D convolutional layers with 32 filters of size 7 and a stride of length 2, and one or two densely connected layers of 256 or 512 units. The output of the last layer is passed through two stacked densely connected layers of hidden size units to evaluate *µ* and *σ* of the re-parameterization trick [35].

**Figure 1:**
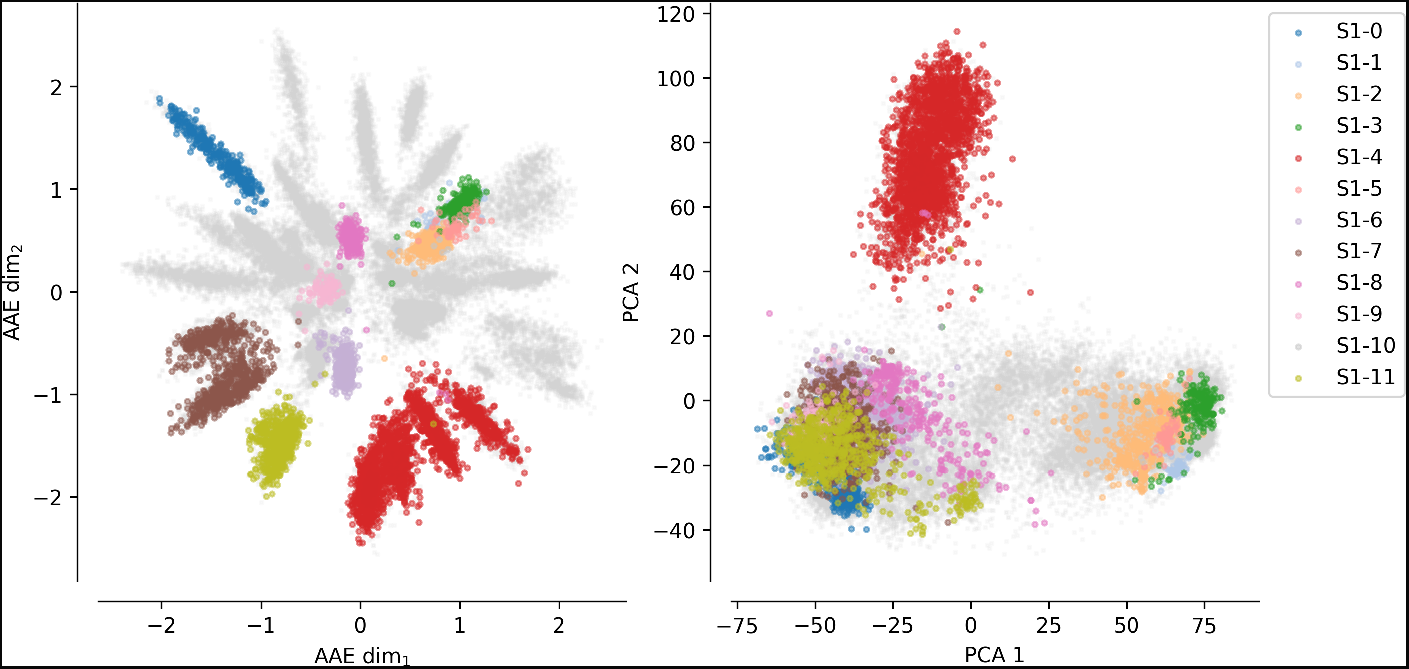
Sequences of the SulfAtlas MSA projected using the encoding learned using an AAE (number of latent dimensions: 2) and a PCA (two first components). Gray data points correspond to protein sequences not part of the curated 12 sub-families.

The decoder is made of two or three densely connected layers of the length of the sequence family time alphabet units for the last layers and of 256 or 512 units for the first or the two first layers. The final output of the decoder is reshaped and a soft-max activation function is applied, corresponding to a probability for each positions associated with each possible amino acids. To convert the probability matrix of the decoder into a sequence, a random sampling according to the probability output was performed at each position. The selected amino acid at a given position is therefore not necessarily the amino acid with the highest probability. The discriminator network is made of two or three densely connected layers, the last layer has only one unit and corresponds to the discriminator classification decision through a sigmoid activation function, the first or the first two layers are made of 256 or 512 units.

The network was trained for each protein family independently. The autoencoder was trained using a categorical cross-entropy loss function between the input data and the predicted sequences by the autoencoder. The discriminator was trained using binary cross-entropy loss function between the input data encoded and the samples from the prior distribution.

### 2.3 Generated sequences and structures analyses

#### Dimensionality reduction

The AAE model can be used to reduce the dimensionality of the sequence space by setting a small latent size. Two dimensionality reductions were tested with latent size of 2 and 100. Latent size of 2 can be easily visualized and a larger latent size of 100 should represent the input data more efficiently as more information can be stored.

#### Clustering

HDBSCAN [36, 37] was used to cluster the sequences in the latent space due to its capacity to handle clusters of different sizes and densities and its performances in high dimensional space. The Euclidean distance metric was used to compute distances between points of the latent space. A minimal cluster size of 60 was set to consider a group as a cluster as the number of protein sequences is rather large. The minimal number of samples in a neighborhood to consider a point as a core point was set to 15 to maintain relatively conservative clusters.

#### Functional and taxonomic analyses

Enzyme functional annotation (EC ids) and NCBI taxonomic identifiers were extracted when available from the Gene Ontology Annotation portal (January 2019) using the UniProt-GOA mapping [38]. Protein without annotation were not taken into account in these analyses.

The annotation homogeneity was computed for each cluster. Considering a cluster, the number of different EC ids and taxonomic ids were retrieved. For each different EC ids (taxonomic ids) its percentage in the cluster was computed. An EC id (taxonomic id) of a cluster with a value of 90% indicates that 90% of the cluster members have this EC id (taxonomic id). A cluster with high homogeneity values correspond to functionally or evolutionary related sequences.

Homogeneous clusters computed from the AAE encoded space will therefore indicates the ability of the AAE model to capture and to distinguish protein sequences with functionally or evolutionary relevant features without supervision.

#### Latent space interpolation

Twenty pairs of protein sequences were randomly chosen between all combinations of sulfatases sub-families with at least 100 labeled members but with less than 1000 members (to avoid pronounced imbalance between classes): S1-0 (308 proteins), S1-2 (462 proteins), S1-3 (186 proteins), S1-7 (741 proteins), S1-8 (290 proteins) and S1-11 (669 proteins). The coordinates of the selected sequences in the encoded space with 100 dimensions were retrieved and spherical interpolation using 50 steps were performed for each of the pairs. Spherical interpolation has previously been reported to provide better interpolation for the generation of images [39]. The interpolated points were given to the decoder and new sequences were generated. Statistical analyses were carried out using the generated sequences based on their dynamical changes from one family to an other to study the model ability to generalize. Analyses at the amino acid level were also performed on the interpolated sequences of two Sulfatase sub-families encoded far from one-another in the latent space.

#### Latent space arithmetic

Subtraction and addition of latent spaces have been shown to be able to transfer features specific to some sub-groups of data to other sub-groups. This property was tested on seven previously selected Sulfatase sub-families (S1-0, S1-1, S1-2, S1-3, S1-7, S1-8 and S1-11) based on their number of protein sequences. Different arithmetic strategies (Supplementary Figure 2) were tested between latent spaces of a query sub-family and a different source sub-family with the aim to transfer features of the source sub-family to the query sub-family.

**Figure 2:**
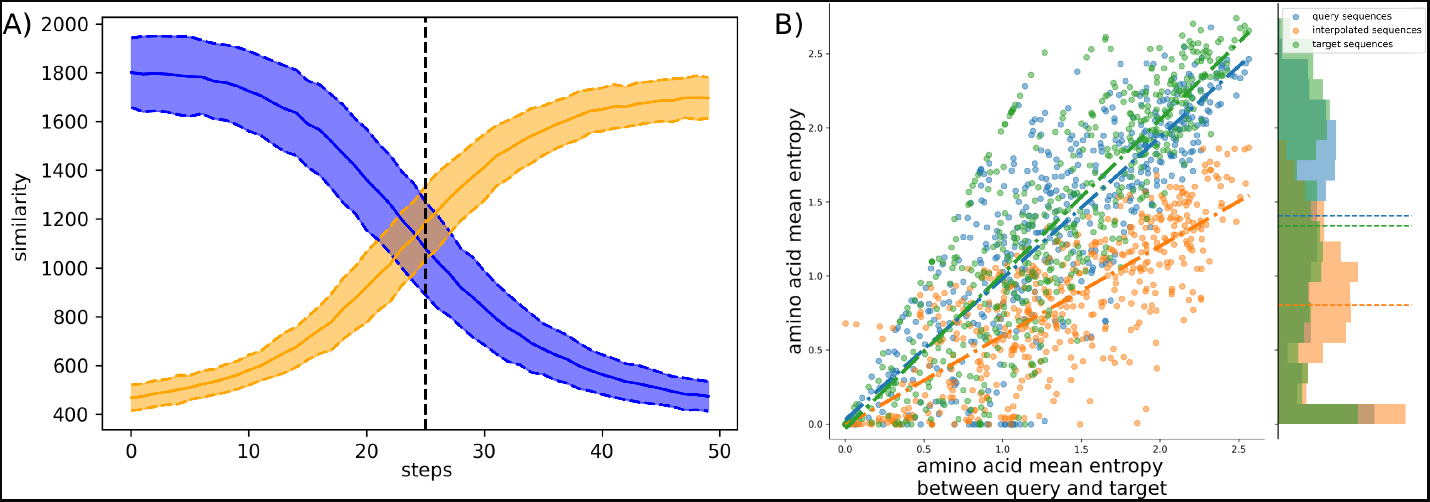
Interpolation analyses between sub-families S1-0 and S1-11. A) Sequence similarity distributions (sum of blosum weights, higher the score higher the similarity) between interpolated sequences and query proteins (blue) or target proteins (orange). B) Distribution of amino acid Shannon entropy for interpolated sequences (orange) between sub-families S1-0 (blue) and S1-11 (green) over the amino acid mean Shannon entropy of query and target sub-families.

A first strategy consists in adding the mean latent space, computed using the encoder on the sequences of the source sub-family, to the encoded sequences of the query sub-family. The second strategy differs from the first one by subtracting the mean background latent space, computed from the latent space of all sub-families, from the latent space of the query sub-family. The third strategy differs from the second as the background strategy is computed using all sub-families except source and query sub-families. Finally, in the fourth strategy, the subtraction is performed using a local KD-tree to only remove features shared by the closest members of a given query and the addition is performed by randomly selecting a member of the source family and its closest 10 members.

For each strategy, new sequences were generated using the latent spaces of all query proteins in the sub-families. Thus, for one original encoded protein sequence there is a direct correspondence between the original amino acid sequence and the generated amino acid sequences with the different strategies and different source sub-families. The generated sequences by latent space arithmetic are compared to the initial sub-families in terms of sequence and structural constraints.

To evaluate the generated sequences by latent space arithmetic, the sequences were compared to themselves and to the biological sequences of the two initial sub-families using protein sequence similarity computed with a Blosum 62 substitution matrix. The sequence similarities inside a sub-family and between sub-families are also computed. The distributions of sequence similarities allow to explore the ability of the latent space arithmetic operations and of the decoder to produce meaningful intermediate protein sequences from unexplored encoded data points.

Protein structural models are computed using the structures of the initial sub-families as template for MODELLER [40] and evaluated using the DOPE score [41]. Models are computed using the generated sequences by latent space arithmetic on template structures of their source and query sub-families. The DOPE energies of the modeled structures are compared to structural models computed as references. The first references are structural models computed using the sequences and template structures of the same sub-families, which should provide the best DOPE energies (best case scenario). The second references are structural models computed using the sequences and template structure of different sub-families (ex: sequences from source sub-family and template structures from the query sub-family or inversely, sequences from query sub-family and template structures from the source sub-family), which should provide the worst DOPE energy (worst case scenario). If the generated sequences by latent space arithmetic correspond to intermediate proteins with properties from two sub-families, they should have intermediate DOPE energies when compared to the best and worst case scenarii.

## 3 Results

A structurally constrained MSA was computed using Expresso from T-Coffee web-server [42, 43] between sequences of S1 sulfatases structures. This MSA was processed into a Hidden Markov Model and hmmsearch was used to retrieve aligned sequence matches against the UniRef90 sequence database. A total of 76,427 protein sequence hits were found to match the sulfatase HMM in UniRef90. The sequences were filtered to remove columns and hits with more than 90% and 75%, respectively, of gap characters. The final MSA comprised 41,901 sequences. The sulfatases protein dataset was separated in training, validation, and test sets with a split ratio of: 0.8, 0.1, and 0.1.

The three different AAE architectures (see Method section) were trained on the training set and evaluated on the validation set. The test set was only used on the final selected architecture. Models were evaluated by computing top k-accuracy, corresponding to the generation of the correct amino acid in the first *k* amino acids. Supplementary Table 2 shows the top k accuracy metric for k=1 and k=3 computed for the different autoencoders. The accuracy scores scaled down with the number of parameters, but without any large difference. To avoid over-fitting, the architecture with the fewest number of parameters (architecture 3) was therefore selected. The final accuracy scores on the test set were computed and were similar to the values observed during the model training: 62.5% and 80.2% (k=1 and k=3). The selected architecture was separately trained on the protein sequences of the HUP and TPP families.

### 3.1 Latent space projection

AAE can be used as a dimensional reduction and visualization techniques by fixing the dimension of the latent space to two or three for plotting purpose. In this section, AAE network ability to create meaningful projection is tested on Sulfatase, HUP and TPP families by clustering and analysing protein sequences in terms of enzymatic activity and phylogenetic diversity.

Starting from the final MSA of the Sulfatase family, an AAE network was trained to project the sequences in a latent space with two dimensions. For comparison, the MSA projection using the first two components of the PCA decomposition was also computed.

Figure 1 shows the protein sequences encoded by the AAE in two latent dimensions and the PCA projections. Each dot corresponds to a protein sequence, and the dots are colored according to their sub-family. Gray dots correspond to protein sequences not belonging to any of the 12 curated sulfatases sub-families. In this figure, the AAE displays a higher capacity to disentangle the sequence and functional spaces of the S1 family than a PCA. Interestingly, it can be observed in the AAE projection some well-separated gray spikes (sequences not belonging to any curated sub-family). These spikes may correspond to groups of enzymes sharing common substrate specificity. In some cases, sub-families with identical functions are projected closely on the encoded space. For instance, sub-families S1-6 (light magenta) and S1-11 (yellow) have both the EC 3.1.6.14 activity (N-acetylglucosamine-6-sulfatase) and are closely located in the encoded space. Moreover, some sub-family projections appear entangled such as the S1-1 sub-family (light blue, Cerebroside sulfatase activity, EC 3.1.6.8), the S1-2 (orange) and the S1-3 (green) sub-families (Steryl-sulfatase activity, EC 3.1.6.2), the S1-5 (pink) sub-family (N-acetylgalactosamine-6-sulfatase activity, EC 3.1.6.4), and the S1-10 (gray) sub-family (Glucosinolates sulfatase activity EC 3.1.6.-). The five families correspond to four different functions but are made of Eukaryotic protein sequences only and their entanglement may be due to their shared common evolutionary history. This separation based on the sequence kingdoms can clearly be visualized in the PCA projections with Eukaryotic sequences on the right side on sub-families with a majority of Bacteria sequences on the left side. The example of protein B6QLZ0 PENMQ is also interesting. The protein is projected (yellow dot corresponding to the S1-11 sub-family) at coordinates (0.733, −1.289), inside the space of the S1-4 family (red). This may look like an error by closer inspection shows that this protein is part of both the S1-4 and S1-11 sub-families of the SulfAtlas database.

Projections of sequences into latent spaces using AAE with two dimensions were also tested on the HUP and TPP families. The AAE projections can be visualized on Supplementary Figure 3. There are fewer functional annotations for these two families, but a strong separation between the major functions of the families can clearly be seen.

**Figure 3:**
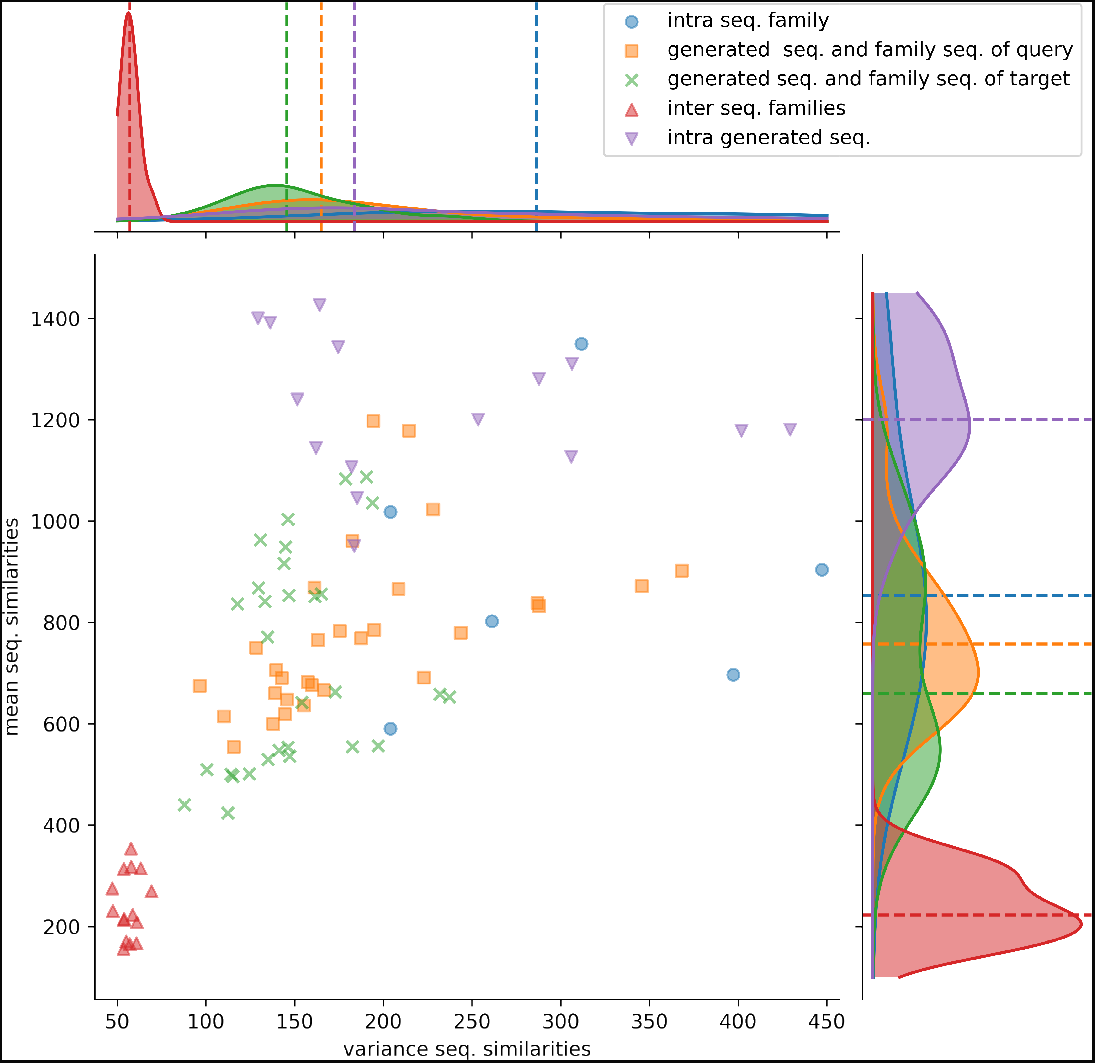
Distributions of protein sequence similarities. Blue dots: protein sequence similarity computed between sequences of the same protein sub-family. Orange squares: similarity computed between generated sequences and the sequences of their query sub-family (ex: S1-0m2 generated sequences and S1-0 sub-family sequences). Green x: similarity computed between generated sequences and the sequences of their target sub-family (ex: S1-0m2 generated sequences and S1-2 sub-family sequences). Red upper triangles: similarity computed between sequences of two different sub-families (ex: S1-0 sequences and S1-2 sequences). Magenta lower triangles: similarity computed between sequences of the same generated sequence group. The variance and the mean of each distribution are displayed on the horizontal and vertical axes.

Latent spaces were evaluated for each protein family based on enzyme classification (EC) and taxonomic homogeneity. Given a set of protein sequences, the encoded sequences in a latent space of a 100 dimensions were clustered using HDBSCAN. For the sulfatase family, 27 clusters were found, for which taxonomic and EC annotations could be extracted (Supplementary Table 3). All these clusters displayed either strong taxonomic or EC homogeneity. Enzymatic homogeneity was higher than taxonomic homogeneity for 16 clusters, found equal in one cluster and lower for 10 clusters. In the HUP family, all clusters had very high EC homogeneity (Supplementary table 4). Only two clusters out of 47 could be found with higher taxonomic homogeneity than EC homogeneity and for this two clusters enzymatic homogeneity values were high and only marginally different (cluster 5, taxonomic homogeneity of 100% an EC homogeneity of 99% and cluster 31, taxonomic homogeneity of 99 % and EC homogeneity of 97%). Five clusters were found with equal taxonomic and EC homogeneity. Similarly, in the TPP family, all clusters had also very high EC homogeneity (Supplementary table 5). Five clusters out of 51 could be found with higher taxonomic homogeneity than EC homogeneity. For these 5 clusters the differences between taxonomic homogeneity and EC homogeneity were higher than the differences observed for the HUP clusters. Six clusters were found with equal taxonomic and EC homogeneity. These results highlight the ability of the encoding space to capture amino acid functional properties instead of signal due to taxonomic relationships.

### 3.2 Protein latent space interpolation

Interpolation between encoded sequences can be used to “navigate” between proteins of two sub-families. For this task, twenty pairs of protein sequences were randomly selected between all combinations of protein sub-families to test the capacity of the encoded space and 50 intermediates data points were generated between each pair. At each step, the sequence similarities were computed between the generated protein sequences from the interpolated latent space and the query and target protein sequences of the sub-families. It is thus possible to measure the amino acid sequence drift from one protein to another one.

The observed amino acid transition from the query sub-family to the target sub-family is very smooth for all combinations of sub-families and has a logistic function shape as shown in Figure 2-A. The smooth transition between points demonstrates the ability of AAE network to encode the sequences in smooth latent space and thus to “fill” the gap between protein sequence sub-families.

The Shannon entropy was computed for each group of sequences: interpolated sequences between query and target sub-families, sequences of the query sub-families and sequences of the target sub-families. Figure 2-B shows the Shannon entropy distribution for the S1-0 and S1-11 sequences and their interpolated sequences. Interestingly, the figure shows lower entropy for interpolated sequences than for original sequences. This trend is true for all interpolated sequences between all sub-families as reported in the Supplementary Table 6. Lower entropy indicates fewer variation in the interpolated amino acids than in biological sequences which point out to restricted paths between sub-families. This is in agreement with molecular evolution theory and experiments that describe protein families as bassins in fitness landscape [44, 45, 46].

A closer inspection of interpolations between sub-families S1-0 and S1-4 (respectively blue and red data points in figure 1) was also performed to study changes at the amino acid level as the two sub-families are in opposite spaces in the two-dimensional projection. It can be observed in Supplementary Figure 5 that gapped area found in the query sequence but not in the target sequence (and inversely) are progressively filled (or broken down) starting from the flanking amino acids to the center of the gap (or inversely from the center to the flanking amino acids). This indicates an organized and progressive accumulation of amino acids (or gaps) that extend (or shrink) the region of the sequence previously without (or with) residues. For instance gap reduction can be observed in the generated sequences between sequence ID 2 of the sulfatase S1-0 family (query) and sequence ID 2196 of the sulfatase S1-4 family (target) at positions 75 to 86.

Moreover, family-specific amino acids are progressively replaced in key positions. In the previous interpolation between query and target sequences, it can thus be observed at positions 21 and 22 of the MSA a replacement of residues S and C by G and A (Supplementary Figure 5). Most transitions are not abrupt, and do not occur at the 25th-generated intermediate sequences but are smooth and correspond to plausible sequences. The ability of the AAE to generate interpolated sequences with emerging or disappearing features of two sub-families, highlights its capacity to generalize the decoding of latent space points not corresponding to encoded sequences and thus never observed during training, and outside the structured organization of the computed latent space.

### 3.3 Protein latent space arithmetic

It has been shown that latent space arithmetic was able to transfer learned features between different classes [30]. If applied to protein sequences, this latent space feature could permit to transfer features such as enzymatic activity or part of structure between protein families. In this work, to test this ability, different arithmetic strategies (see Methods and Supplementary Figure 2) were tested between latent spaces of two Sulfatase sub-families with the aim to transfer features from a source sub-family to a query sub-family.After performing the arithmetic operation between latent space coordinates, the protein sequences corresponding to the new coordinates were generated by the decoder. The sequences were then modelled on protein structure templates corresponding to both sub-families and compared to generated models using sequences and structures of the same sub-families and using sequences from one sub-family and structure templates from the other one. The Sulfatases sub-families S1-0, S1-2, S1-3, S1-7, S1-8 and S1-11 were chosen to test this hypothesis.

In the following section, the terminology Seq. S1-XmY will correspond to a generated sequence using a combination of latent space of the sub-family S1-Y added to the latent space of the sub-family S1-X. The X and Y sub-families will be referred to as the query and source sub-families. The source latent space is obtained using the mean value of the latent space corresponding to the sequences of the sub-family

First, two Prosite motifs of the Sulfatase family are analyzed from generated and original sequences. Supplementary Figure 6 displays logo plots of two regions corresponding to Prosite motifs PS00523 and PS00149 to illustrate the amino acid content of the generated protein sequences by latent space arithmetic. These regions correspond to the most conserved regions of the sulfatase family and have been proposed as signature patterns for all the sulfatases in the Prosite database.

Different amino acid patterns can be observed between the sequence groups that can be classified as “competition”, “taking over”, or “balanced” pattern. A competition pattern of amino acids corresponds to equivalent frequency of two different amino acids in the generated sequences. A taking over pattern corresponds to an amino of one of the original sequences being the most frequent in the generated sequences. Finally, a balanced pattern corresponds to a maintained equilibrium between amino acids in the generated sequences. Other positions are however displaying much more complex patterns and cannot be summarized as a frequency competition between source and query sub-families. These behaviors can be observed several times through the logo plots but are still position-specific, meaning that the bits scores pattern observed in the source sub-families (Panels A and D) do not necessary allow to predict the amino acids bits scores in the generated sequences (Panels B and C).

Protein sequence similarities were computed to evaluate the diversity of the generated sequences and to compare their diversity with the original sub-families. Protein sequence similarities were computed between:

- sequences of a group of generated sequences,
- sequences of a sulfatase sub-family used to generate protein sequences,
- generated sequences and their query sulfatase sub-family,
- generated sequences and their source sulfatase sub-family,
- query and source sequences of sulfatase sub-families.

Figure 3 shows the mean and variance distribution of computed protein sequence similarities between these different groups for generated sequences computed using the first strategy. The first, second, and third strategies display a similar pattern and their corresponding figures are available in the Supplementary Information (Supplementary Figures 10, 12 and 14).

Protein sequence similarities between different sub-families (red upper triangles) have lower similarity scores and lower variances than the other distributions. Protein sequence similarities between sequences of a sub-family (blue circles) have the highest mean and variance values observed. However, since only 6 sub-families were kept for analysis (sub-families 0, 2, 3, 7, 8, and 11), trends must therefore be taken with precaution. Generated protein sequences compared to themselves (magenta lower triangles) have mean and variance protein sequence similarities higher than when compared to their query or sub-families. The last two have (generated sequences compared to query sequences, orange squares and generated sequences compared to target sequences, green crosses) mean and variance values spread between the blue and red distributions.

These distributions indicate that generated protein sequences by latent space arithmetic have an intrinsic diversity similar to the biological sub-families. Moreover, the generated sequences are less similar to the sequences from their query and source sub-families than to themselves. The generated sequences are also globally as similar to the sequences of their query sub-family as to the sequence of their source sub-family. The generation process is therefore able to capture the features of the selected query and source sub-families and to generate a protein sequence diversity similar to the original sub-families.

Finally, protein structure modeling was performed to assess and compare the properties of the generated sequences by latent space arithmetic and the protein sequences of the natural sub-families. For each sub-family, 100 original sequences were randomly selected along the corresponding generated sequences. All the generated sequences were aligned to protein structures of their corresponding source and query sub-families, and the alignments were used to create 3D structures models by comparative modeling. The quality of models was then evaluated with the DOPE function of MODELLER.

Supplementary Figure 7 shows an example of the energy distribution computed from models using the second strategy with query sub-family S1-0 and source sub-family S1-2. The lowest energies (best models) were found on modelled structures using the original protein sequences of a sub-family to the structural templates of the same sub-family (*Struct. 0 Seq. 0* and *Struct. 2 Seq. 2*). Conversely, the highest energies are found on modelled structures using the original protein sequences of a sub-family to the structural templates of another sub-family (*Struct. 0 Seq. 2* and *Struct. 2 Seq. 0*). Interestingly, generated sequences using additions and subtractions of latent spaces have intermediate energy distributions. This can be clearly observed in Figure 4, where generated sequences are mostly situated between the two dotted lines. Dots on the right side of the vertical line at 0 correspond to modeled structures using sequences of the latent space with lower energy than the modeled structures using sequences from their original sub-family. Dots on the left side of the vertical line at 0 are modeled structures using sequences of the latent space with higher energy than the modeled structures using sequences from their original sub-family. The diagonal line on the top-left corner corresponds to the difference in energy between modeled structures using sequences from their original sub-family and modeled structures using sequences from biological sequences of another sub-family. Generated sequences modeled on structures belonging to the same sub-family than their query latent space sub-family (ex: *Struct. 0 Seq.S1-0m2* and *Struct. 2 Seq. S1-2m0* on Supplementary Figure 7 and *M_QS_/Q* on Figure 4) have slightly lower energy than when modeled on structures corresponding to the sub-family of their source latent space sub-family (ex: *Struct. 0 Seq. S1-2m0* and *Struct. 2 Seq. S1-0m2* on Supplementary Figure 7 and *M_SQ_/Q* on Figure 4). This trend is true for all query / source pairs of sub-families and all strategies except for generated sequences using the fourth strategy (local background subtraction of query latent space using a KD-tree and the addition of source latent space), see Supplementary Figures 9, 11, 13 and Methods.

**Figure 4:**
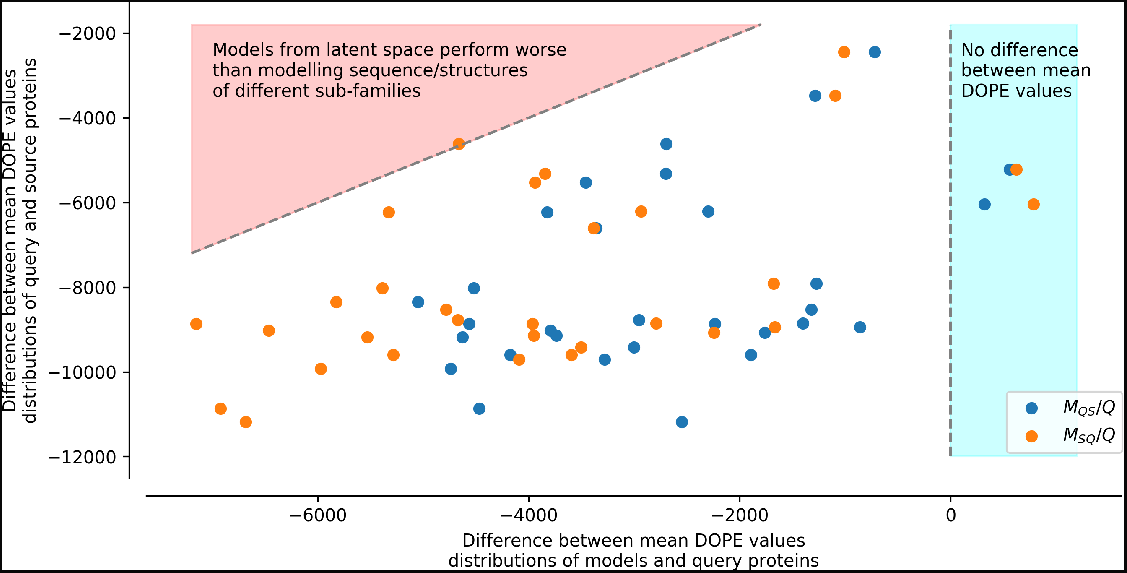
Difference between mean DOPE distributions. Mean value for each distribution, such as the distributions presented in Figure Supplementary 7, were computed. The *y* axis represents the difference between the mean values computed for query sequences modeled on structures of the same sub-family and mean values computed for source sequences modeled on structures of the query sub-family (ex: differences between mean of Struct. 0 Seq. 0 and mean of Struct. 0 Seq. 2 distributions in Supplementary Figure 7). The *x* axis corresponds to the difference between the mean values computed for query sequences modeled on structures of the same sub-family and mean values computed for query sequences to which latent spaces of the source sub-family sequences have been added and modeled on structures of the query sub-family (*M_QS_ /Q*), or source sequences to which latent spaces of the query sub-family sequences have been added and modeled on structures of the source sub-family (*M_SQ_/Q*) (ex: differences between mean of Struct. 0 Seq. S1-0m2 and mean of Struct. 0 Seq. 0 distributions Supplmementary in Figure 7).Points in the red area correspond to mean distribution values from generated sequences whose modeled structures have a higher energy than models created using pairs of sequences/structures from different sub-families. Points in the blue area correspond to mean distribution values from generated sequences whose modeled structures have a lower energy than models created using pairs of sequences/structures from the same sub-family.

In this strategy, the modeled structures using generated sequences do not display energy distributions in-between the energy distributions of the original sequences modeled on structures of the query or of the source sub-families (dotted lines). The energy distribution of generated sequences modeled on structures belonging to the sub-family of their query latent space sub-family (ex: *Struct. 0 Seq.S1-0m2*, blue dots *M_QS_/Q*) with the fourth strategy is closer to the energy distribution of the modeled structures using a sequence and a structure template from the same sub-families. The energy distribution of generated sequences modeled on structures corresponding to the sub-family of their source latent space (ex: *Struct. 2 Seq.S1-0m2*, orange dots *M_SQ_/Q*) with the fourth strategy is closer to the energy distribution of the modeled structures using a sequence and a structure template from different sub-families. This indicates that the fourth strategy is less robust to latent space arithmetic than the other three strategies. No clear differences could be observed between the first, second, and third strategy.

## 4 Discussion

In this study, a new framework is presented to analyze and explore the protein sequence space regarding functionality and evolution. Previous works based on Variational Autoencoder (VAE) have successfully reported the ability of this deep learning framework to model protein sequence and functional spaces [16], predict amino acid fitness impact [17, 18], look into protein evolution [19] or design new protein [20]. In this study, an other unsupervised framework based on Adversarial Autoencoder (AAE) is proposed to disentangle the information contained in protein sequences and to capture functional and evolutionary specific features. AAE networks have the advantage over VAEs to constrain the latent space over a prior distribution which has previously been reported to have better efficiency[22]. After checking the ability of AAE networks to disentangle correctly protein functional space, this study proposes to explore the capacity of AAE networks with two sampling tasks: interpolate protein sequences of different sub-families and generate new protein sequences mixing properties of two sub-families.

First, AAEs were trained on protein sequence families known to have different sub-families with specific functions. The results highlight the ability of AAEs to separate sequences according to functional and taxonomic properties for the three studied families. This emphasized the ability of the AAEs to extract biological relevant features and encode them accordingly to their biological differences into the latent space. Second, interpolations between encoded sequences in the latent space of different sub-families show smooth transitions regarding amino acid substitutions even at the active site positions. The generated sequences along the interpolation paths are intermediate sequences with properties similar to their closest sub-families forming the start or end points of the path. The generated sequences have amino acid Shannon entropy lower than sequences from sub-families which indicates a, amino acid diversity at each position lower than the biological sequences. This trend can be expected as the interpolation method takes the shortest path between the two points of the sub-families. Finally, three strategies were explored to generate protein sequences with features from two different sub-families. These strategies are based on latent space arithmetic, a concept previously applied in image generation tasks to produce relevant images with intermediate features. Three out of the four different experiments carried out were able to generate sequences with intermediate features as measured from their protein sequence similarity distributions and modeling energy assessments. Biological experiments will be needed to confirm the functional relevance of the transferred features, but the strategies could have many applications should it be validated.

The absence of measured differences between three out of four strategies used to generate intermediate sequences may also indicate that more optimal approaches could be designed. In this regard, the model architecture could also be improved. Currently, the model input is a filtered MSA, but improved architectures could probably benefit from full protein sequences of different sizes without filtering. It is for instance known that some motifs in the TPP and HUP family plays important roles in the family sub-functions [6]. As protein specific motifs, they are not necessary conserved and may not reach filtering thresholds. Recent advances have been made regarding the protein sequence universe representation notably using self-sueprvised approaches and Transformer models [47, 48, 49, 50, 51]. The reported techniques in this study can be applied to any latent space projection and it would be interesting to combine them with representation of the protein sequence universe to navigate and perform feature transfer between protein families.

The results of this study show that AAE, in particular, and deep learning generative models in general, can provide original solutions for protein design and functional exploration.

## 5 Acknowledgements

TBF’s work was suported by ISCD, Sorbonne Université, Paris, France. TBF thanks the NVIDIA society for providing a TitanXp GPU to perform computations.

## Notes

### Competing Interest Statement

The authors have declared no competing interest.

### Summary of Updates

Update Acknowledgement

## References

[1] Axe DD. Estimating the prevalence of protein sequences adopting functional enzyme folds. Journal of molecular biology. 2004;341(5):1295–1315.

[2] Luisi PL, Chiarabelli C, Stano P. From never born proteins to minimal living cells: two projects in synthetic biology. Origins of Life and Evolution of Biospheres. 2006;36(5):605–616.

[3] Dawson NL, Lewis TE, Das S, Lees JG, Lee D, Ashford P, et al. CATH: an expanded resource to predict protein function through structure and sequence. Nucleic acids research. 2016;45(D1):D289–D295.

[4] El-Gebali S, Mistry J, Bateman A, Eddy SR, Luciani A, Potter SC, et al. The Pfam protein families database in 2019. Nucleic acids research. 2018;47(D1):D427–D432.

[5] Pandurangan AP, Stahlhacke J, Oates ME, Smithers B, Gough J. The SUPERFAMILY 2.0 database: a significant proteome update and a new webserver. Nucleic acids research. 2018;47(D1):D490–D494.

[6] Das S, Dawson NL, Orengo CA. Diversity in protein domain superfamilies. Curr Opin Genet Dev. 2015 Dec;35:40–49.

[7] Tian P, Louis JM, Baber JL, Aniana A, Best RB. Co-Evolutionary Fitness Landscapes for Sequence Design. Angewandte Chemie International Edition. 2018;57(20):5674–5678.

[8] Copp JN, Akiva E, Babbitt PC, Tokuriki N. Revealing unexplored sequence-function space using sequence similarity networks. Biochemistry. 2018;57(31):4651–4662.

[9] Salinas VH, Ranganathan R. Coevolution-based inference of amino acid interactions underlying protein function. elife. 2018;7:e34300.

[10] Tubiana J, Cocco S, Monasson R. Learning protein constitutive motifs from sequence data. elife. 2019;8:e39397.

[11] Poelwijk FJ, Socolich M, Ranganathan R. Learning the pattern of epistasis linking genotype and phenotype in a protein. Nature communications. 2019;10(1):1–11.

[12] Russ WP, Figliuzzi M, Stocker C, Barrat-Charlaix P, Socolich M, Kast P, et al. An evolution-based model for designing chorismate mutase enzymes. Science. 2020;369(6502):440–445.

[13] Lopez R, Regier J, Cole MB, Jordan MI, Yosef N. Deep generative modeling for single-cell transcriptomics. Nature methods. 2018;15(12):1053.

[14] Gómez-Bombarelli R, Wei JN, Duvenaud D, Hernández-Lobato JM, Sánchez-Lengeling B, Sheberla D, et al. Automatic Chemical Design Using a Data-Driven Continuous Representation of Molecules. ACS Central Science. 2018;4(2):268–276. Available from: https://doi.org/10.1021/acscentsci.7b00572.

[15] Rampasek L, Hidru D, Smirnov P, Haibe-Kains B, Goldenberg A. Dr.VAE: Drug Response Variational Autoencoder. arXiv e-prints. 2017 Jun;p. arXiv:1706.08203.

[16] Sinai S, Kelsic E, Church GM, Nowak MA. Variational auto-encoding of protein sequences. arXiv preprint arXiv:171203346. 2017;.

[17] Riesselman AJ, Ingraham JB, Marks DS. Deep generative models of genetic variation capture the effects of mutations. Nat Methods. 2018;15:816–822.

[18] Hopf TA, Ingraham JB, Poelwijk FJ, Schärfe CP, Springer M, Sander C, et al. Mutation effects predicted from sequence co-variation. Nature biotechnology. 2017;35(2):128–135.

[19] Ding X, Zou Z, Brooks III CL. Deciphering protein evolution and fitness landscapes with latent space models. Nature communications. 2019;10(1):1–13.

[20] Greener JG, Moffat L, Jones DT. Design of metalloproteins and novel protein folds using variational autoencoders. Scientific reports. 2018;8(1):1–12.

[21] Makhzani A, Shlens J, Jaitly N, Goodfellow IJ. Adversarial Autoencoders. CoRR. 2015;abs/1511.05644. Available from: http://arxiv.org/abs/1511.05644.

[22] Kadurin A, Nikolenko S, Khrabrov K, Aliper A, Zhavoronkov A. druGAN: an advanced generative adversarial autoencoder model for de novo generation of new molecules with desired molecular properties in silico. Molecular pharmaceutics. 2017;14(9):3098–3104.

[23] Goodfellow I, Pouget-Abadie J, Mirza M, Xu B, Warde-Farley D, Ozair S, et al. Generative adversarial nets. Advances in neural information processing systems. 2014;27:2672–2680.

[24] Barbeyron T, Brillet-Gueguen L, Carre W, Carriere C, Caron C, Czjzek M, et al. Matching the Diversity of Sulfated Biomolecules: Creation of a Classification Database for Sulfatases Reflecting Their Substrate Specificity. PLoS ONE. 2016;11(10):e0164846.

[25] Bork P, Koonin EV. AP-loop-like motif in a widespread ATP pyrophosphatase domain: implications for the evolution of sequence motifs and enzyme activity. Proteins: Structure, Function, and Bioinformatics. 1994;20(4):347–355.

[26] Wolf YI, Aravind L, Grishin NV, Koonin EV. Evolution of aminoacyl-tRNA synthetases—analysis of unique domain architectures and phylogenetic trees reveals a complex history of horizontal gene transfer events. Genome research. 1999;9(8):689–710.

[27] Aravind L, Anantharaman V, Koonin EV. Monophyly of class I aminoacyl tRNA synthetase, USPA, ETFP, photolyase, and PP-ATPase nucleotide-binding domains: implications for protein evolution in the RNA world. Proteins: Structure, Function, and Bioinformatics. 2002;48(1):1–14.

[28] Muller YA, Lindqvist Y, Furey W, Schulz GE, Jordan F, Schneider G. A thiamin diphosphate binding fold revealed by comparison of the crystal structures of transketolase, pyruvate oxidase and pyruvate decarboxylase. Structure. 1993;1(2):95–103.

[29] Berthold CL, Moussatche P, Richards NG, Lindqvist Y. Structural basis for activation of the thiamin diphosphate-dependent enzyme oxalyl-CoA decarboxylase by adenosine diphosphate. Journal of Biological Chemistry. 2005;280(50):41645–41654.

[30] Radford A, Metz L, Chintala S. Unsupervised representation learning with deep convolutional generative adversarial networks. arXiv preprint arXiv:151106434. 2015;.

[31] Suzek BE, Wang Y, Huang H, McGarvey PB, Wu CH, Consortium U. UniRef clusters: a comprehensive and scalable alternative for improving sequence similarity searches. Bioinformatics. 2014;31(6):926–932.

[32] Eddy SR. Accelerated profile HMM searches. PLoS computational biology. 2011;7(10):e1002195.

[33] Orengo CA, Michie A, Jones S, Jones DT, Swindells M, Thornton JM. CATH–a hierarchic classification of protein domain structures. Structure. 1997;5(8):1093–1109.

[34] Sillitoe I, Dawson N, Lewis TE, Das S, Lees JG, Ashford P, et al. CATH: expanding the horizons of structure-based functional annotations for genome sequences. Nucleic acids research. 2018;47(D1):D280–D284.

[35] Kingma DP, Welling M. Auto-encoding variational bayes. Oral presentation at the International Conference on Learning Representations, Banff, Alberta, Canada. 2014 14–16 April;.

[36] Campello RJ, Moulavi D, Sander J. Density-based clustering based on hierarchical density estimates. In: Pacific-Asia conference on knowledge discovery and data mining. Springer; 2013. p. 160–172.

[37] McInnes L, Healy J. Accelerated Hierarchical Density Based Clustering. In: Data Mining Workshops (ICDMW), 2017 IEEE International Conference on. IEEE; 2017. p. 33–42.

[38] Huntley RP, Sawford T, Mutowo-Meullenet P, Shypitsyna A, Bonilla C, Martin MJ, et al. The GOA database: gene ontology annotation updates for 2015. Nucleic acids research. 2014;43(D1):D1057–D1063.

[39] White T. Sampling Generative Networks: Notes on a Few Effective Techniques. CoRR. 2016;abs/1609.04468. Available from: http://arxiv.org/abs/1609.04468.

[40] Webb B, Sali A. Comparative protein structure modeling using MODELLER. Current protocols in bioinformatics. 2014;47(1):5–6.

[41] Shen MY, Sali A. Statistical potential for assessment and prediction of protein structures. Protein Sci. 2006 Nov;15(11):2507–2524.

[42] Armougom F, Moretti S, Poirot O, Audic S, Dumas P, Schaeli B, et al. Expresso: automatic incorporation of structural information in multiple sequence alignments using 3D-Coffee. Nucleic Acids Res. 2006 Jul;34(Web Server issue):W604–608.

[43] Di Tommaso P, Moretti S, Xenarios I, Orobitg M, Montanyola A, Chang JM, et al. T-Coffee: a web server for the multiple sequence alignment of protein and RNA sequences using structural information and homology extension. Nucleic Acids Res. 2011 Jul;39(Web Server issue):W13–17.

[44] Bornberg-Bauer E, Chan HS. Modeling evolutionary landscapes: mutational stability, topology, and superfunnels in sequence space. Proceedings of the National Academy of Sciences. 1999;96(19):10689–10694.

[45] Sikosek T, Chan HS, Bornberg-Bauer E. Escape from adaptive conflict follows from weak functional trade-offs and mutational robustness. Proceedings of the National Academy of Sciences. 2012;109(37):14888–14893.

[46] Boucher JI, Cote P, Flynn J, Jiang L, Laban A, Mishra P, et al. Viewing protein fitness landscapes through a next-gen lens. Genetics. 2014;198(2):461–471.

[47] Rives A, Goyal S, Meier J, Guo D, Ott M, Zitnick CL, et al. Biological Structure and Function Emerge from Scaling Unsupervised Learning to 250 Million Protein Sequences. bioRxiv. 2019;Available from: https://www.biorxiv.org/content/early/2019/04/29/622803.

[48] Alley EC, Khimulya G, Biswas S, AlQuraishi M, Church GM. Unified rational protein engineering with sequence-based deep representation learning. Nature methods. 2019;16(12):1315–1322.

[49] Heinzinger M, Elnaggar A, Wang Y, Dallago C, Nechaev D, Matthes F, et al. Modeling aspects of the language of life through transfer-learning protein sequences. BMC bioinformatics. 2019;20(1):1–17.

[50] Rao R, Bhattacharya N, Thomas N, Duan Y, Chen X, Canny J, et al. Evaluating protein transfer learning with tape. Advances in Neural Information Processing Systems. 2019;32:9689.

[51] Strodthoff N, Wagner P, Wenzel M, Samek W. UDSMProt: universal deep sequence models for protein classification. Bioinformatics. 2020;36(8):2401–2409.

