## Supplementary Information for "Navigating the amino acid sequence space between functional proteins using a deep learning framework"

March 24, 2021

### 1 Supplementary Information

#### 1.1 Deep Neural Network architecture

Supplementary Figure 1 presents the Deep Neural Network architecture used in the project and Supplementary Table 1 the different network architecture variations.

Table 1: Differences between layers of the evaluated model architectures

| Architecture | 1 | 2 | 3 |
|---|---|---|---|
| Encoder | Conv 1D (32, 7)<br>Dense (512) | 2 x Conv 1D (32, 7)<br>2 x Dense (512) | 2 x Conv 1D (32, 7)<br>2 x Dense (256 |
| Decoder | Dense (512) | 2 x Dense (512) | 2 x Dense (256) |
| Discriminator | Dense (512) | 2 x Dense (512) | 2 x Dense (256) |

Table 2 reports the top-k (k=1 and k=3) accuracy metrics for the three architectures described in Supplementary Table 1 on the Training and validation data sets.

Table 2: Accuracy metrics (k=1 and k=3) on the train and validation sets of sulfatases using different models (see Method)

| Architecture | k=1 accuracy (%) | | k=3 accuracy (%) | | Number of |
|---|---|---|---|---|---|
| | Train | Validation | Train | Validation | Parameters |
| 1 | 67.3 | 66 | 83.9 | 82.3 | 11 198 553 |
| 2 | 65.1 | 64 | 82.4 | 81 | 9 769 593 |
| 3 | 63.3 | 62.4 | 81.2 | 80.1 | 8 073 337 |

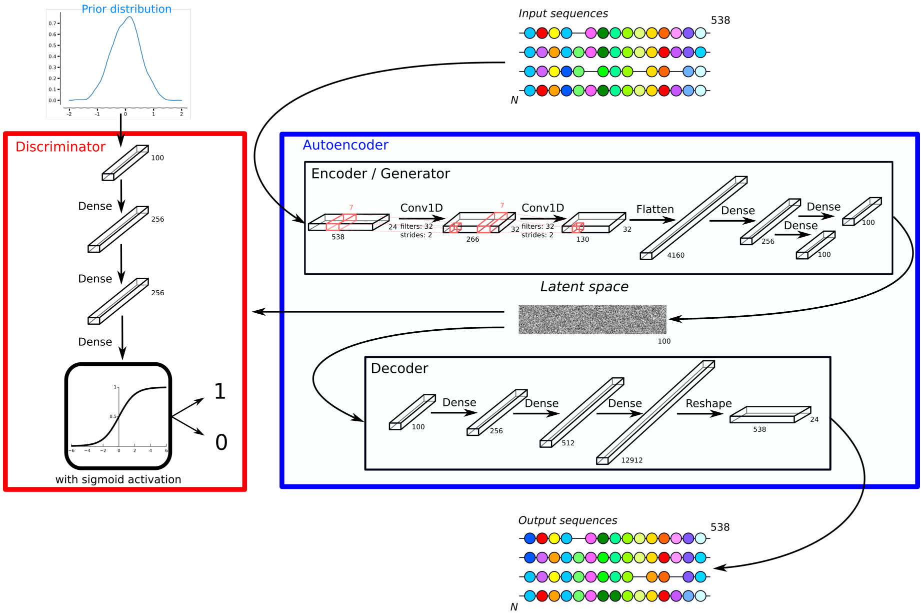


Figure 1: The Adversarial AutoEncoder architecture number 3 presented in Supplementary Table 1. The discriminator (in red) takes as input data from a prior distribution or the latent space computed by the encoder/generator. Using a sigmoid activation function, the discriminator is trained to distinguish between the two types of data. By updating the weight of the encoder/generator based on the discriminator performances, the encoder/generator learn to approximate the prior distribution and fool the discriminator. The autoencoder architecture (in blue) corresponds to a variational autoencoder. Latent space is decoded by a decoder and new sequences are generated using a softmax activation function.

### 1.2 Structural modeling pipeline

Using the latent space computed by the neural network described in Supplementary Figure 1, four different latent space arithmetic strategies were developed to produce sequences with mixed functions. The generated protein sequences were further modelled using *MODELLER* and their energy evaluated using the *DOPE* energy function. Supplementary Figure 2 describes the general pipeline developed.

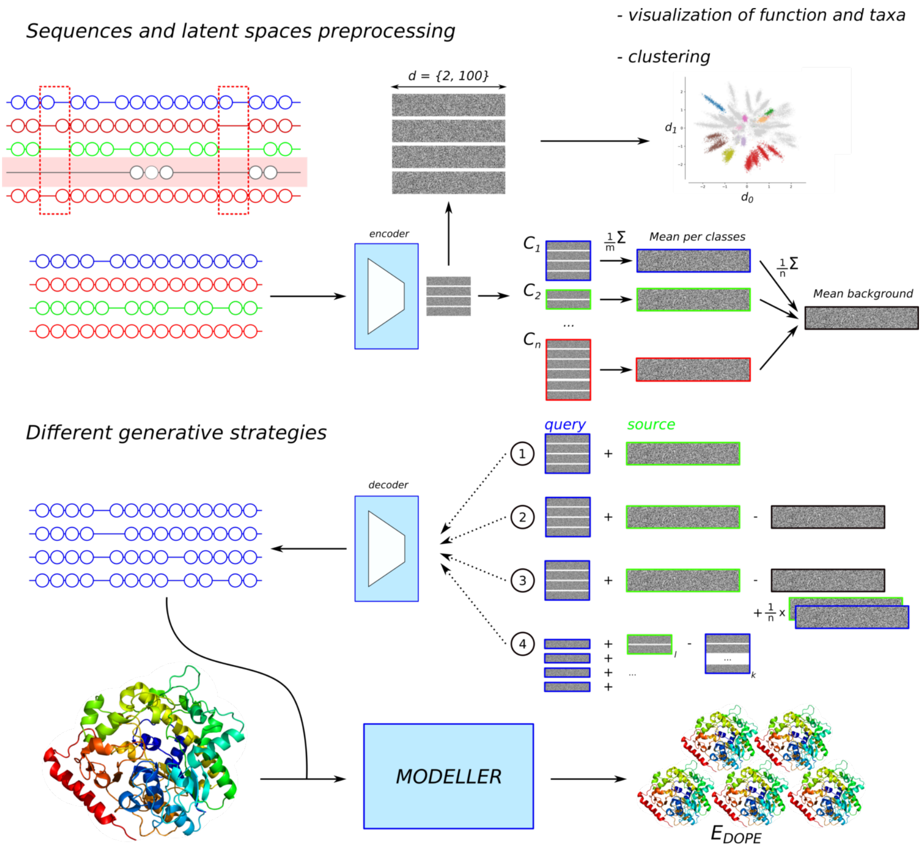


Figure 2: Modeling pipeline used to generate sequences sharing properties of two sub-families. The *hmmsearch* MSA is filtered and passed to the encoder to project each sequence to the latent space. Latent space projections can be used for visualization (see Supplementary Figure 4). Different strategies (1 to 4) are tested to generate new points in the latent space and generate new sequences through the decoder. The new sequences are used in combination with structures of the sub-families to create homology-based structural models and evaluated using the *DOPE* energy function of *MODELLER*.

### 1.3 HUP and TPP latent spaces

Latent spaces generated by the neural network described Supplementary Figure 1 were evaluated in terms of functional and taxonomic properties for the Sulfatase, HUP and TPP families (see main text for the results regarding the Sulfatase family).

Supplementary Figure 3 presents the latent spaces of the HUP and TPP families colored according to their EC annotations. HUP points colored in yellow correspond to protein with EC 6.1.1.1 and 6.1.1.2, pink-colored points to proteins with EC 6.1.1.1 and violet colored points to proteins with EC 2.7.11.24 and 6.1.1.2. TPP sequences have more annotated functions than HUP sequences (57 different EC assignation), but a global pattern can be found in the projection corresponding to two groups of proteins (brown and violet) annotated with EC 2.2.1.1 (Transketolase), oxidoreductase proteins (EC 1.-.-.-, in orange, pink, red, and green shades), and proteins with function EC 2.2.1.9 (2-succinyl-5-enolpyruvyl-6-hydroxy-3-cyclohexene-1-carboxylic-acid synthase, in yellow and gray shades).

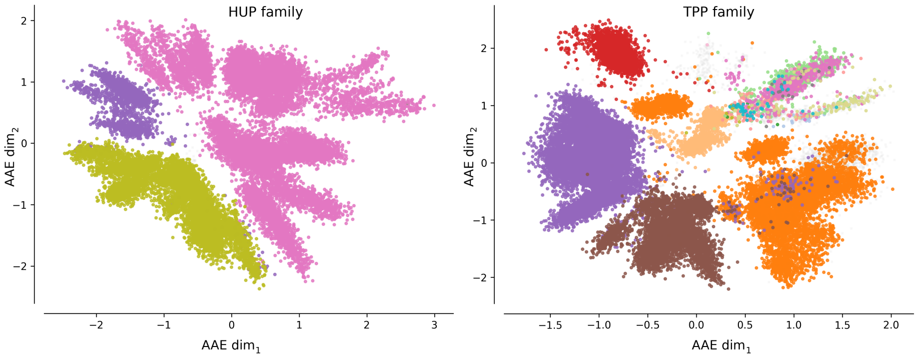


Figure 3: Encoded sequences of HUP (left) and TPP (right) multiple sequences alignments using AAEs with 2 latent dimensions. Data points are colored according to their enzyme classification annotations retrieved from GOA.

### 1.4 Enzymatic and taxonomic specificity

Left Supplementary Figure 4 displays a clustering results in a 100 latent dimensional projected into a 2 latent dimensional latent space. GOA Enzyme Classification annotation are used to color the right plot.

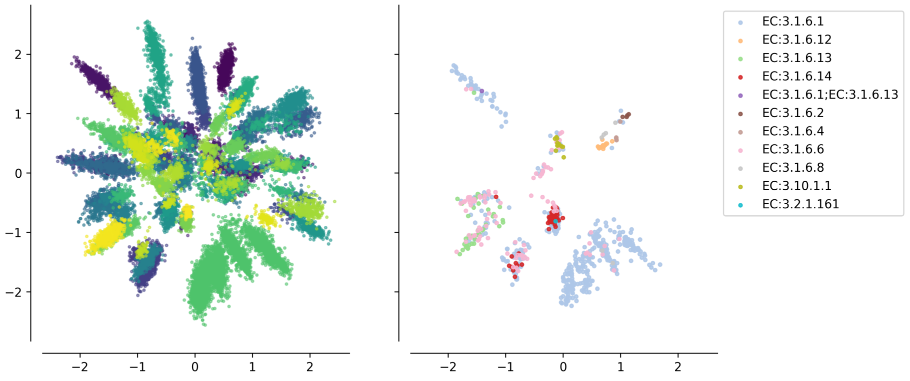


Figure 4: Projection of SulfAtlas MSA encoded sequences using an AAE with two latent dimension. Left - sequences are colored based on the clustering of results on protein sequences projected into an AAE with 100 latent dimensions. Right - sequences are colored according to the retrieved GOA Enzyme Classification annotation.

Supplementary Tables 3 displays for each cluster computed in the 100 dimensional latent space its homogeneity in term of taxonomy and functional annotation. Clusters have a higher functional homogeneity than taxonomic homogeneity. Thus, sequences are projected according to sequence signal coming from functional activity rather than evolutionary relationship.

Table 3: Enzymatic classes and taxonomic homogeneity of encoded sulfatases after clustering by HDBSCAN.

| cluster index | Taxonomic group | percentage of proteins with identical taxa | Enzyme class | percentage of proteins with identical EC |
|---|---|---|---|---|
| 12 | Proteobacteria | 0.34 | EC:3.1.6.1 | 0.96 |
| 27 | Proteobacteria | 0.98 | EC:3.1.6.1 | 0.89 |
| 28 | Proteobacteria | 0.93 | EC:3.1.6.1 | 1.00 |
| 34 | Bacteroidetes | 0.65 | EC:3.1.6.6 | 0.67 |
| 38 | Ascomycota | 0.63 | EC:3.10.1.1 | 0.50 |
| 39 | Arthropoda | 0.48 | EC:3.10.1.1 | 1.00 |
| 44 | Ascomycota | 0.91 | EC:3.1.6.1 | 1.00 |
| 45 | Chordata | 0.60 | EC:3.1.6.14 | 1.00 |
| 46 | Actinobacteria | 1.00 | EC:3.1.6.14 | 0.67 |
| 47 | Bacteroidetes | 0.69 | EC:3.1.6.6 | 0.69 |
| 63 | Bacteroidetes | 0.67 | EC:3.1.6.1 | 0.60 |
| 64 | Bacteroidetes | 0.81 | EC:3.1.6.1 | 0.70 |
| 65 | Bacteroidetes | 0.64 | EC:3.1.6.14 | 0.50 |
| 66 | Bacteroidetes | 0.69 | EC:3.1.6.1 | 1.00 |
| 67 | Ascomycota | 0.49 | EC:3.1.6.1 | 0.50 |
| 68 | Bacteroidetes | 0.95 | EC:3.1.6.6 | 0.67 |
| 73 | Planctomycetes | 0.46 | EC:3.1.6.6 | 0.71 |
| 74 | Bacteroidetes | 0.80 | EC:3.1.6.6 | 0.50 |
| 76 | Planctomycetes | 0.51 | EC:3.1.6.6 | 0.67 |
| 84 | Chordata | 0.42 | EC:3.1.6.13 | 1.00 |
| 86 | Bacteroidetes | 0.62 | EC:3.1.6.13 | 0.40 |
| 87 | Bacteroidetes | 0.74 | EC:3.1.6.13 | 0.83 |
| 101 | Chordata | 0.88 | EC:3.1.6.2 | 0.44 |
| 102 | Chordata | 0.80 | EC:3.1.6.4 | 1.00 |
| 103 | Chordata | 0.89 | EC:3.1.6.8 | 1.00 |
| 111 | Chordata | 0.82 | EC:3.1.6.12 | 1.00 |
| 112 | Arthropoda | 0.97 | EC:3.1.6.12 | 1.00 |

Supplementary Tables 4 and 5 show the taxonomic and functional homogeneity of clusters for the HUP and TPP families with similar findings.

Table 4: Enzymatic classes and taxonomic homogeneity of encoded HUP proteins after clustering by HDBSCAN.

| cluster index | Taxonomic group | percentage of proteins with identical taxa | Enzyme class | percentage of proteins with identical EC |
|---|---|---|---|---|
| 0 | Candidatus | 0.70 | EC:6.1.1.1 | 1.00 |
| 1 | Candidatus | 0.74 | EC:6.1.1.1 | 1.00 |
| 2 | Candidatus | 0.38 | EC:6.1.1.1 | 1.00 |
| 3 | Euryarchaeota | 0.98 | EC:6.1.1.2 | 1.00 |
| 4 | Euryarchaeota | 0.96 | EC:6.1.1.1 | 1.00 |
| 5 | Streptophyta | 1.00 | EC:6.1.1.1 | 0.99 |
| 6 | Chloroflexi | 0.90 | EC:6.1.1.1 | 1.00 |
| 7 | Arthropoda | 0.98 | EC:6.1.1.1 | 1.00 |
| 8 | Ascomycota | 0.99 | EC:6.1.1.1 | 1.00 |
| 9 | Crenarchaeota | 0.46 | EC:6.1.1.1 | 1.00 |
| 10 | Chordata | 1.00 | EC:6.1.1.1 | 1.00 |
| 11 | Euryarchaeota | 0.94 | EC:6.1.1.1 | 1.00 |
| 12 | Streptophyta | 0.36 | EC:6.1.1.1 | 1.00 |
| 13 | Euryarchaeota | 0.24 | EC:6.1.1.2 | 1.00 |
| 14 | Ascomycota | 0.32 | EC:6.1.1.2 | 0.94 |
| 15 | Bacteroidetes | 1.00 | EC:6.1.1.1 | 1.00 |
| 16 | Candidatus | 0.35 | EC:6.1.1.1 | 1.00 |
| 17 | Ascomycota | 0.41 | EC:6.1.1.1 | 1.00 |
| 18 | Euryarchaeota | 0.96 | EC:6.1.1.2 | 1.00 |
| 19 | Ascomycota | 0.99 | EC:6.1.1.2 | 1.00 |
| 20 | Candidatus | 0.86 | EC:6.1.1.2 | 1.00 |
| 21 | Chloroflexi | 0.28 | EC:6.1.1.2 | 1.00 |
| 22 | Proteobacteria | 0.92 | EC:6.1.1.1 | 1.00 |
| 23 | Candidatus | 0.71 | EC:6.1.1.2 | 1.00 |
| 24 | Cyanobacteria | 0.98 | EC:6.1.1.1 | 1.00 |
| 25 | Proteobacteria | 0.99 | EC:6.1.1.1 | 1.00 |
| 26 | Firmicutes | 0.96 | EC:6.1.1.1 | 1.00 |
| 27 | Firmicutes | 0.98 | EC:6.1.1.1 | 1.00 |
| 28 | Proteobacteria | 0.98 | EC:6.1.1.1 | 1.00 |
| 29 | Firmicutes | 1.00 | EC:6.1.1.1 | 1.00 |
| 30 | Firmicutes | 0.98 | EC:6.1.1.1 | 1.00 |
| 31 | Actinobacteria | 0.99 | EC:6.1.1.2 | 0.97 |
| 32 | Chordata | 0.86 | EC:6.1.1.2 | 1.00 |
| 33 | Bacteroidetes | 0.96 | EC:6.1.1.1 | 1.00 |
| 34 | Firmicutes | 0.98 | EC:6.1.1.1 | 1.00 |
| 35 | Actinobacteria | 0.99 | EC:6.1.1.1 | 1.00 |
| 36 | Actinobacteria | 1.00 | EC:6.1.1.1 | 1.00 |
| 37 | Proteobacteria | 0.99 | EC:6.1.1.1 | 1.00 |
| 38 | Proteobacteria | 0.99 | EC:6.1.1.1 | 1.00 |
| 39 | Proteobacteria | 0.56 | EC:6.1.1.2 | 1.00 |
| 40 | Proteobacteria | 0.99 | EC:6.1.1.2 | 1.00 |
| 41 | Firmicutes | 0.73 | EC:6.1.1.2 | 1.00 |
| 42 | Proteobacteria | 0.99 | EC:6.1.1.2 | 1.00 |
| 43 | Bacteroidetes | 0.97 | EC:6.1.1.2 | 1.00 |
| 44 | Actinobacteria | 1.00 | EC:6.1.1.2 | 1.00 |
| 45 | Actinobacteria | 1.00 | EC:6.1.1.2 | 1.00 |
| 46 | Proteobacteria | 0.96 | EC:6.1.1.2 | 1.00 |

Table 5: Enzymatic classes and taxonomic homogeneity of encoded TPP proteins after clustering by HDBSCAN.

| cluster index | Taxonomic group | percentage of proteins with identical taxa | Enzyme class | percentage of proteins with identical EC |
|---|---|---|---|---|
| 0 | Firmicutes | 0.75 | EC:2.2.1.1 | 1.00 |
| 1 | Bacteroidetes | 1.00 | EC:2.2.1.1 | 1.00 |
| 2 | Thermotogae | 0.32 | EC:2.2.1.1 | 1.00 |
| 3 | Ascomycota | 0.73 | EC:1.2.4.1 | 1.00 |
| 4 | Proteobacteria | 0.44 | EC:1.2.7.3 | 1.00 |
| 5 | Proteobacteria | 0.84 | EC:1.2.4.4 | 1.00 |
| 6 | Proteobacteria | 0.67 | EC:2.2.1.1 | 0.96 |
| 7 | Proteobacteria | 0.73 | EC:2.2.1.1 | 1.00 |
| 8 | Proteobacteria | 0.90 | EC:2.2.1.7 | 1.00 |
| 9 | Actinobacteria | 0.82 | EC:1.2.4.1 | 0.68 |
| 10 | Candidatus | 0.80 | EC:2.2.1.1 | 1.00 |
| 11 | Firmicutes | 0.67 | EC:2.2.1.7 | 1.00 |
| 12 | Proteobacteria | 0.64 | EC:2.2.1.1 | 1.00 |
| 13 | Firmicutes | 0.54 | EC:2.2.1.1 | 1.00 |
| 14 | Firmicutes | 0.89 | EC:2.2.1.1 | 0.98 |
| 15 | Euryarchaeota | 0.91 | EC:2.2.1.1 | 1.00 |
| 16 | Actinobacteria | 0.44 | EC:2.2.1.1 | 0.99 |
| 17 | Proteobacteria | 0.74 | EC:1.2.4.1 | 1.00 |
| 18 | Actinobacteria | 0.68 | EC:1.2.3.3 | 0.33 |
| 19 | Proteobacteria | 0.57 | EC:1.2.7.1 | 1.00 |
| 20 | Verrucomicrobia | 0.77 | EC:2.2.1.7 | 1.00 |
| 21 | Candidatus | 0.46 | EC:2.2.1.1 | 1.00 |
| 22 | Bacteroidetes | 0.97 | EC:2.2.1.1 | 0.86 |
| 23 | Firmicutes | 0.62 | EC:3.7.1.22 | 0.95 |
| 24 | Verrucomicrobia | 0.88 | EC:2.2.1.1 | 1.00 |
| 25 | Proteobacteria | 0.97 | EC:3.7.1.22 | 1.00 |
| 26 | Proteobacteria | 1.00 | EC:3.7.1.22 | 1.00 |
| 27 | Firmicutes | 1.00 | EC:2.2.1.7 | 1.00 |
| 28 | Proteobacteria | 0.98 | EC:1.2.4.1 | 1.00 |
| 29 | Actinobacteria | 0.87 | EC:1.2.4.1 | 1.00 |
| 30 | Proteobacteria | 0.93 | EC:2.2.1.1 | 0.99 |
| 31 | Firmicutes | 0.98 | EC:2.2.1.7 | 1.00 |
| 32 | Cyanobacteria | 0.33 | EC:1.2.4.1 | 1.00 |
| 33 | Ascomycota | 0.99 | EC:2.2.1.3 | 0.50 |
| 34 | Actinobacteria | 0.92 | EC:3.7.1.22 | 0.72 |
| 35 | Firmicutes | 0.99 | EC:2.2.1.7 | 1.00 |
| 36 | Proteobacteria | 0.83 | EC:2.2.1.7 | 1.00 |
| 37 | Bacteroidetes | 0.98 | EC:2.2.1.7 | 1.00 |
| 38 | Firmicutes | 0.35 | EC:1.2.7.1 | 0.68 |
| 39 | Bacteroidetes | 0.97 | EC:2.2.1.7 | 1.00 |
| 40 | Bacteroidetes | 0.98 | EC:2.2.1.7 | 1.00 |
| 41 | Actinobacteria | 0.99 | EC:2.2.1.7 | 1.00 |
| 42 | Actinobacteria | 0.98 | EC:2.2.1.7 | 1.00 |
| 43 | Actinobacteria | 0.99 | EC:2.2.1.6 | 0.99 |
| 44 | Proteobacteria | 0.93 | EC:2.2.1.6 | 1.00 |
| 45 | Cyanobacteria | 1.00 | EC:2.2.1.6 | 1.00 |
| 46 | Proteobacteria | 0.88 | EC:2.2.1.6 | 1.00 |
| 47 | Proteobacteria | 0.98 | EC:2.2.1.6 | 1.00 |
| 48 | Ascomycota | 1.00 | EC:2.2.1.6 | 1.00 |
| 49 | Proteobacteria | 0.58 | EC:2.2.1.1 | 1.00 |
| 50 | Proteobacteria | 0.58 | EC:2.2.1.1 | 0.92 |

### 1.5 Protein latent space interpolation

Supplementary Figure 5 shows the sequences generated during the interpolation between protein sequences of S1-0 and S1-4 sub-families .

An abrupt transition can be observed at positions 53, G (S1-0 sub-family) to S (S1-4 sub-family), and 51 [VI] (S1-0 sub-family) to T (S1-4 sub-family), corresponding to very conserved residues in Prosite motif PS00523 (see Supplementary Figure 6). The other positions of the motif are less affected by abrupt transitions but also appear to have lower amino acid diversity than other columns. A similar behavior can only be observed for position 111, T (S1-0 sub-family) to W (S1-4 sub-family), of motif PS00149 (positions 102 to 112). The other positions of the motifs are either conserved (T105, G109, K110) or accepting fluctuations (columns 103, 104, 107).

Supplementary Table 6 displays the interpolation coefficients along Perason correlation and $R^2$ values of interpolations between pairs of sub-families (S1-0, S1-2, S1-3, S1-6, S1-7, S1-8 and S1-11). The computed coefficients correspond to the linear regression presented in the main document Figure 2.

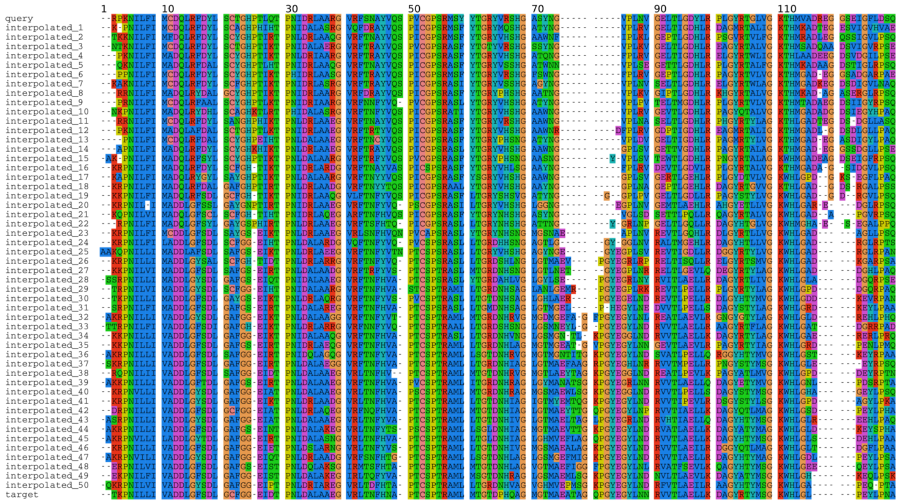

Figure 5: **First 130 amino acids for generated sequences using interpolation between data points of the latent space.** The interpolation is performed between latent spaces of protein ID 2 of the sulfatase S1-0 family and of protein ID 2196 of the sulfatase S1-4 sub-family. Amino acid color coding is based on physo-chemical properties. Large transitions between gaps to amino acids and amino acids to gaps can be observed at positions 75 to 86 and positions 116 to 122. Amino acid columns transformation can be observed at multiple positions: 21 (S to G), 51 (V/I to T/S), 53 (G to S) etc.

Table 6: Fitted coefficients and correlations between computed amino acid Shannon entropy for original (query/target) and interpolated protein sequence groups.

| query sub-family | target sub-family | computed on | slope | intercept | Pearson correlation | $R^2$ |
|---|---|---|---|---|---|---|
| 0 | 2 | query | 0.956290 | -0.103755 | 0.809619 | 0.655484 |
| 0 | 2 | interpolated | 0.548000 | 0.112658 | 0.691604 | 0.478316 |
| 0 | 2 | target | 1.043710 | 0.103755 | 0.832977 | 0.693850 |
| 0 | 3 | query | 1.031314 | 0.085408 | 0.805843 | 0.649383 |
| 0 | 3 | interpolated | 0.650507 | 0.100268 | 0.758494 | 0.575313 |
| 0 | 3 | target | 0.968686 | -0.085408 | 0.787623 | 0.620349 |
| 0 | 6 | query | 0.993397 | -0.212751 | 0.875553 | 0.766592 |
| 0 | 6 | interpolated | 0.597583 | -0.022404 | 0.788073 | 0.621059 |
| 0 | 6 | target | 1.006603 | 0.212751 | 0.878228 | 0.771285 |
| 0 | 7 | query | 0.953058 | -0.186997 | 0.890607 | 0.793182 |
| 0 | 7 | interpolated | 0.637465 | -0.120335 | 0.845739 | 0.715274 |
| 0 | 7 | target | 1.046942 | 0.186997 | 0.906816 | 0.822316 |
| 0 | 8 | query | 1.068595 | -0.245278 | 0.860925 | 0.741193 |
| 0 | 8 | interpolated | 0.621773 | -0.024649 | 0.799847 | 0.639755 |
| 0 | 8 | target | 0.931405 | 0.245278 | 0.827715 | 0.685112 |
| 0 | 11 | query | 0.956111 | 0.029197 | 0.852433 | 0.726643 |
| 0 | 11 | interpolated | 0.605286 | -0.006790 | 0.809397 | 0.655124 |
| 0 | 11 | target | 1.043889 | -0.029197 | 0.871847 | 0.760117 |
| 2 | 3 | query | 1.065313 | 0.202054 | 0.882602 | 0.778986 |
| 2 | 3 | interpolated | 0.685519 | -0.055323 | 0.710578 | 0.504922 |
| 2 | 3 | target | 0.934687 | -0.202054 | 0.854806 | 0.730694 |
| 2 | 6 | query | 1.031616 | -0.108499 | 0.885286 | 0.783732 |
| 2 | 6 | interpolated | 0.544565 | 0.048658 | 0.696711 | 0.485406 |
| 2 | 6 | target | 0.968384 | 0.108499 | 0.872652 | 0.761522 |
| 2 | 7 | query | 0.989150 | -0.073795 | 0.879910 | 0.774242 |
| 2 | 7 | interpolated | 0.546516 | 0.050054 | 0.714461 | 0.510455 |
| 2 | 7 | target | 1.010850 | 0.073795 | 0.884159 | 0.781738 |
| 2 | 8 | query | 1.104741 | -0.147410 | 0.886245 | 0.785429 |
| 2 | 8 | interpolated | 0.559824 | -0.002677 | 0.652964 | 0.426363 |
| 2 | 8 | target | 0.895259 | 0.147410 | 0.840368 | 0.706218 |
| 2 | 11 | query | 0.995848 | 0.144879 | 0.868813 | 0.754836 |
| 2 | 11 | interpolated | 0.540960 | 0.086570 | 0.704842 | 0.496802 |
| 2 | 11 | target | 1.004152 | -0.144879 | 0.870572 | 0.757896 |
| 3 | 6 | query | 0.962288 | -0.292272 | 0.802938 | 0.644709 |
| 3 | 6 | interpolated | 0.628164 | 0.071557 | 0.742000 | 0.550565 |
| 3 | 6 | target | 1.037712 | 0.292272 | 0.823697 | 0.678477 |
| 3 | 7 | query | 0.916941 | -0.263226 | 0.811708 | 0.658869 |
| 3 | 7 | interpolated | 0.639068 | 0.038983 | 0.770087 | 0.593034 |
| 3 | 7 | target | 1.083059 | 0.263226 | 0.854012 | 0.729337 |
| 3 | 8 | query | 1.042807 | -0.324905 | 0.835960 | 0.698829 |
| 3 | 8 | interpolated | 0.698894 | -0.053498 | 0.761837 | 0.580396 |
| 3 | 8 | target | 0.957193 | 0.324905 | 0.813383 | 0.661592 |
| 3 | 11 | query | 0.917803 | -0.052419 | 0.774132 | 0.599281 |
| 3 | 11 | interpolated | 0.622037 | 0.115285 | 0.735195 | 0.540512 |
| 3 | 11 | target | 1.082197 | 0.052419 | 0.821733 | 0.675244 |
| 6 | 7 | query | 0.961538 | 0.031202 | 0.919089 | 0.844724 |
| 6 | 7 | interpolated | 0.530712 | -0.018224 | 0.710257 | 0.504465 |
| 6 | 7 | target | 1.038462 | -0.031202 | 0.929441 | 0.863861 |
| 6 | 8 | query | 1.067627 | -0.036331 | 0.905576 | 0.820069 |
| 6 | 8 | interpolated | 0.605831 | -0.095900 | 0.725207 | 0.525926 |
| 6 | 8 | target | 0.932373 | 0.036331 | 0.881242 | 0.776588 |
| 6 | 11 | query | 0.966571 | 0.245659 | 0.902724 | 0.814911 |
| 6 | 11 | interpolated | 0.565156 | -0.057655 | 0.728569 | 0.530812 |
| 6 | 11 | target | 1.033429 | -0.245659 | 0.913369 | 0.834244 |
| 7 | 8 | query | 1.106311 | -0.067482 | 0.919691 | 0.845831 |
| 7 | 8 | interpolated | 0.607767 | -0.109265 | 0.787255 | 0.619770 |
| 7 | 8 | target | 0.893689 | 0.067482 | 0.884120 | 0.781668 |
| 7 | 11 | query | 1.006478 | 0.220907 | 0.905675 | 0.820248 |
| 7 | 11 | interpolated | 0.560187 | -0.026458 | 0.759299 | 0.576535 |
| 7 | 11 | target | 0.993522 | -0.220907 | 0.903546 | 0.816396 |
| 8 | 11 | query | 0.896201 | 0.270295 | 0.864645 | 0.747611 |
| 8 | 11 | interpolated | 0.570043 | -0.053888 | 0.709186 | 0.502945 |
| [illegible] | [illegible] | [illegible] | [illegible] | [illegible] | [illegible] | [illegible] |

### 1.6 Latent space arithmetic

#### 1.6.1 Logo plots

Logo plots were computed on Prosite motifs PS00523 and PS00149 of the Sulfatase family to illustrate the amino acid content of the protein sequences generated by latent space arithmetic . These regions correspond to the most conserved regions of the Sulfatase family and have been proposed as signature patterns for all the sulfatases in the Prosite database.

In Supplementary Figure 6, panels A and D correspond to the biological sequences of the S1-0 sub-family and of the S1-2 sub-family respectively. Panels B and C correspond to generated protein sequences using either the Sulfatase sub-family S1-0 as the source and S1-2 as the query (Panel B) and to which the background latent space has been subtracted (strategy 2 on Supplementary Figure 2) and reciprocally (Panel C).

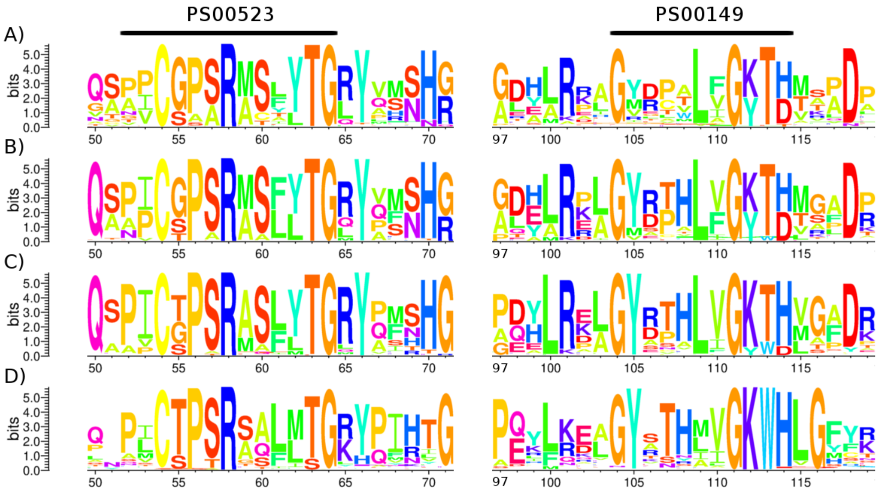


Figure 6: **Logo plots of MSA parts from the S1-0 and S1-2 (panels A and D) sub-families, and generated sequences using S1-0 as query and S1-2 as source (panel B) and S1-2 as query and S1-0 as the source (panel C).**

In the first fragment corresponding to motif PS00523, G55 and T55 are the most frequent amino acid of sub-families S1-0 and S1-2 (Panels A and D) and it can be observed a "competition" between these two amino acids for generated sequences (Panel B and C) with a slightly higher probability for the amino acid of the family used as a query (G in Panel B and T in panel C). The residue S57, involved in the active site of the sulfatases, is less frequent in the query sub-family S1-0 (panel A) than in the sub-family S1-2 (panel D). The high frequency of S at position 57 in the sub-family S1-2 compared to the sub-family S1-1 may have an impact when performing the latent space arithmetic as S is predominant in the generated sequences. This influence of a very frequent amino acid in one of the source or query sub-family on the generated sequences can also be observed at the position 70 and is less visible when multiple amino acid frequencies are more balanced. In the second fragment corresponding to motif PS00149, residue

R at position 101 follows this pattern. It is highly frequent in sub-family S1-0 (panel A) and less frequent in the sub-family S1-2 (panel D). The generated protein sequences display a R at position 101 with high frequency. The inverse can be observed for Y at position 105, highly frequent in sub-family S1-2 (panel D).

Other positions are however displaying much more complex patterns and cannot be summarized as a frequency competition between source and query sub-families. For instance, G at position 71 is very frequent in the sub-family S1-2 but have a comparable frequency with R in sub-family S1-0. The generated protein sequences don't display G has the only possible residue but seem to follow the frequency of their respective query sub-families. Positions in generated sequences with multiple amino acid sharing comparable frequencies in the source and query sub-families have also mixed frequencies, such as in positions 53, 59, 67, 98, or 106.

These behaviors can be observed several times through the logo plots but are still positions specifics, meaning that the bits scores pattern observed in the source sub-families (Panels A and D) do not necessary allow to predict the amino acids bits scores in the generated sub-families (Panels B and C). For instance, W at positions 113 as a high bit score value in the MSA of the sub-family S1-2 but little influences in the amino acids of the generated sequences where T of the sub-family S1-0 is found predominantly. Moreover, these observations are performed on residues of the Prosite motifs which are by definition conserved in sulfatases.

#### 1.6.2 Energy distribution example

Supplementary Figure 7 shows the DOPE energy distributions of generated and original protein sequences with different protein structure templates.

#### 1.6.3 Strategy 1

First strategy consists to add the mean latent space, computed using the encoder on the sequences of the source sub-family, to the encoded sequences of the query sub-family. Supplementary Figure 8 shows the energy distribution

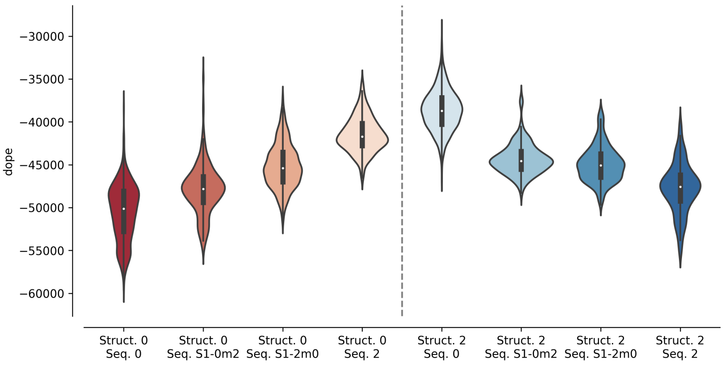


Figure 7: Energy distributions of models computed using structures from sub-family S1-0 (reds) or sub-family S1-2 (blues) and sequences from biological proteins or inferred using latent space arithmetic between spaces encoded by the S1-0 and S1-2 sub-families. Each violin plot corresponds to a specific targeted structure and sequence couples. For example, *Struct. 0 Seq. 0* indicates that the energy distribution corresponds to sequences of the S1-0 sub-family modeled on structures of the S1-0 sub-families and *Struct. 2 Seq. S1-0m2* corresponds to the energy distribution of sequences inferred using the latent space of the sub-family S1-2 added to the latent space of the sub-family S1-0 and modeled on structures of the S1-2 sub-family.

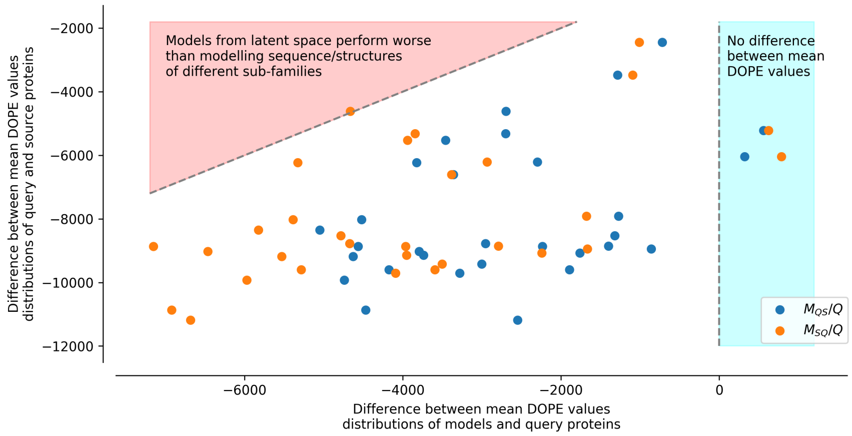


Figure 8: Difference between mean DOPE distributions. Mean value for each distribution, such as the distributions presented in Figure 7, were computed. The $y$ axis represents the difference between the mean values computed for query sequences modeled on structures of the same sub-family and mean values computed for source sequences modeled on structures of the query sub-family (ex: differences between mean of Struct. 0 Seq. 0 and mean of Struct. 0 Seq. 2 distributions in Figure 7). The $x$ axis corresponds to the difference between the mean values computed for query sequences modeled on structures of the same sub-family and mean values computed for query sequences to which latent space of the source sub-family sequences have been added and modeled on structures of the query sub-family ($M_{QS}/Q$), or source sequences to which latent space of the query sub-family sequences have been added and modeled on structures of the source sub-family ($M_{SQ}/Q$) (ex: differences between mean of Struct. 0 Seq. S1-0m2 and mean of Struct. 0 Seq. 0 distributions in Figure 7). Points in the red area correspond to mean distribution values from generated sequences whose modeled structures have a higher energy than models created using pairs of sequences/structures from different sub-families. Points in the blue area correspond to mean distribution values from generated sequences whose modeled structures have a lower energy than models created using pairs of sequences/structures from the same sub-family.

#### 1.6.4 Strategy 2

Second tested strategy differs from the first one by subtracting the mean background latent space, computed from the latent space of all sub-families, from the latent space of the query sub-family.

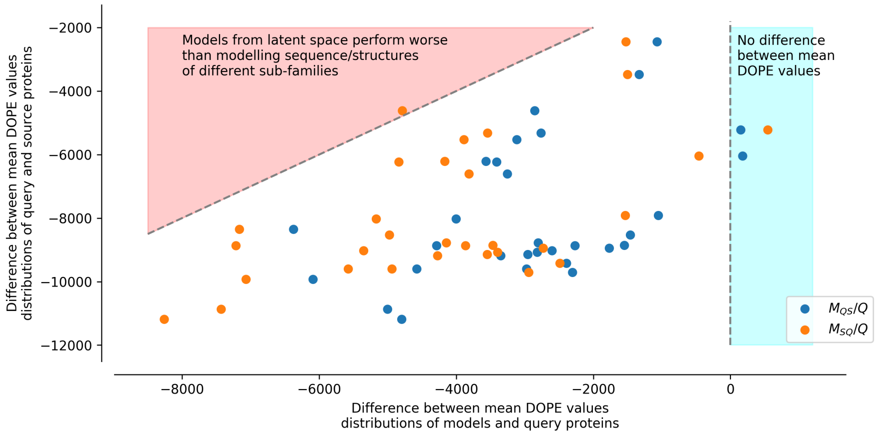


Figure 9: See Figure 8 for legend.

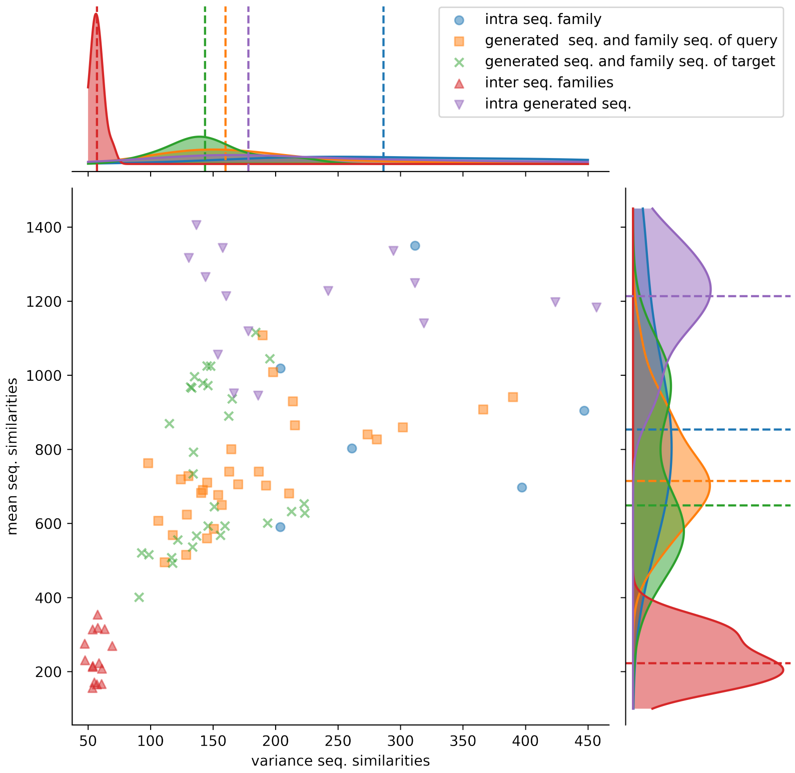


Figure 10: **Distributions of protein sequence similarities**. Blue dots correspond to protein sequence similarity computed between sequences of the same protein sub-family. Orange squares to similarity computed between generated sequences and the sequences of their query sub-family (ex: S1-0m2 generated sequences and S1-0 sub-family sequences). Green x to similarity computed between generated sequences and the sequences of their target sub-family (ex: S1-0m2 generated sequences and S1-2 sub-family sequences). Red upper triangles to similarity computed between sequences of two different sub-families (ex: S1-0 sequences and S1-2 sequences). Magenta lower triangles to similarity computed between sequences of the same generated sequence group. The variance and the mean of each distribution are displayed on the horizontal and vertical axes.

#### 1.6.5 Strategy 3

Third strategy differs to the second as the background strategy is computed using all sub-families except sub-families source and query.

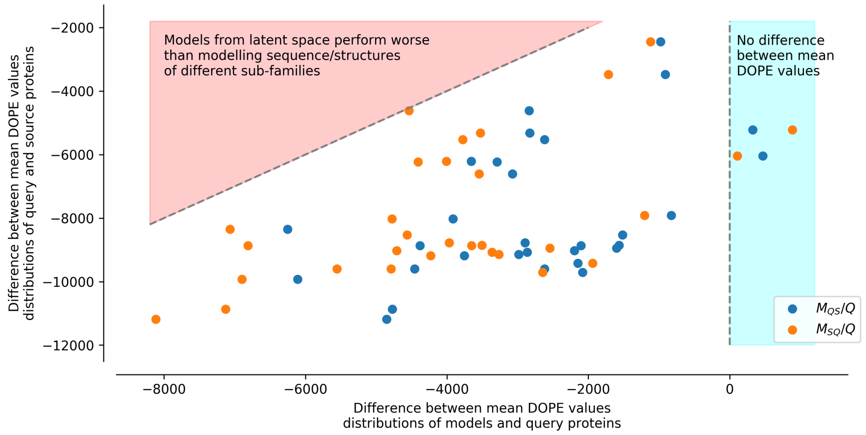


Figure 11: See Figure 8 for legend.

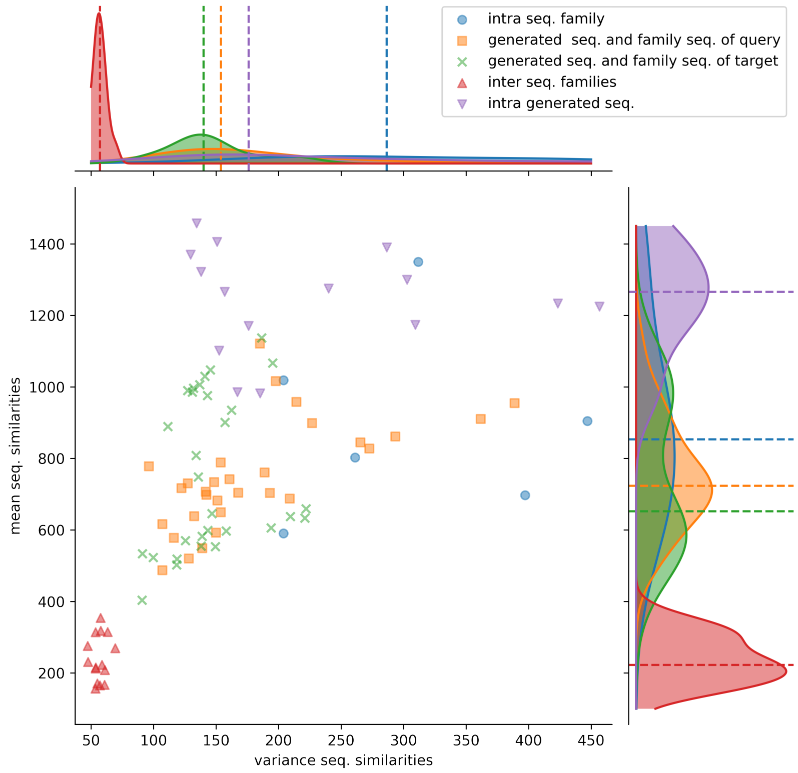


Figure 12: See Figure 10 for legend.

#### 1.6.6 Strategy 4

In the fourth strategy, the subtraction is performed using a local KD-tree to only remove features shared by closest members of a given query and addition is performed randomly selecting a member of the the source family and it's closest 10 members.

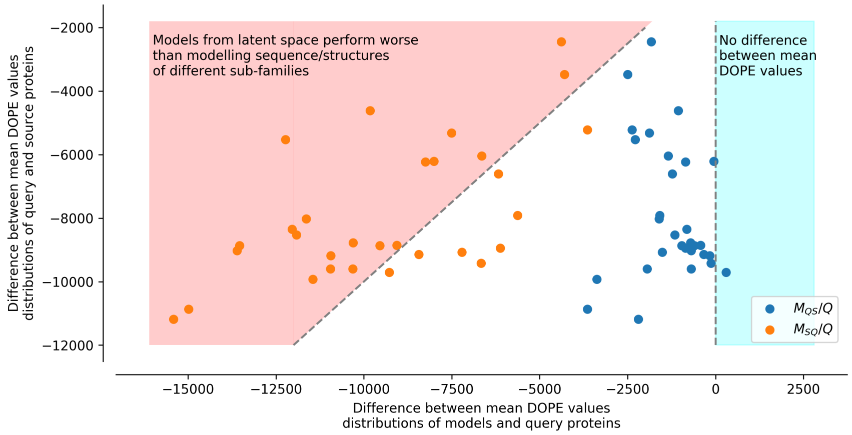


Figure 13: See Figure 8 for legend.

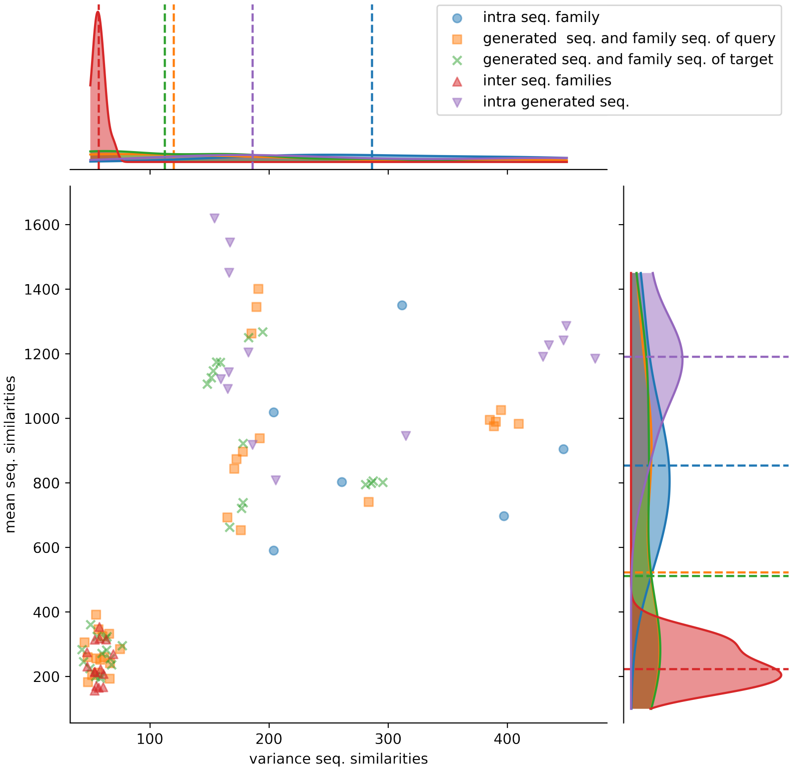


Figure 14: See Figure 10 for legend.
